# HPCA1 is required for systemic ROS and calcium cell-to-cell signaling and plant acclimation to stress

**DOI:** 10.1101/2022.03.24.485694

**Authors:** Yosef Fichman, Sara I Zandalinas, Scott Peck, Sheng Luan, Ron Mittler

## Abstract

As multicellular organisms, plants constantly balance and coordinate many metabolic, physiological, and molecular responses between different cell types and tissues. This process is essential for plant development, growth, and response to different environmental cues. Because plants lack a nervous system, they transmit different signals over long distances via cell-to-cell signaling. Recent studies revealed that reactive oxygen species (ROS), produced by respiratory burst oxidase homologs (RBOHs) at the apoplast play a key role in cell-to-cell signaling. A state of enhanced ROS production by one cell is thereby sensed by a neighboring cell, causing it to produce ROS, creating a continuous chain of cell-to-cell ROS accumulation termed the ‘ROS wave’. This process was found to mediate systemic signals throughout the plant and is required for plant acclimation to different stresses. Although RBOHs were found to produce ROS essential for this process, the identity of the receptor(s) perceiving the apoplastic ROS signal is currently unknow. Here we reveal that the leucine-rich-repeat receptor-like kinase HPCA1 (H_2_O_2_-induced Ca^2+^ increases 1) acts as a central ROS receptor required for the propagation of cell-to-cell ROS signals, systemic signaling in response to different biotic and abiotic stresses, and plant acclimation to stress. We further report that HPCA1 is required for systemic calcium signals, but not systemic membrane depolarization responses, and identify key calcium-dependent signal transduction proteins involved in this process. Our findings reveal that HPCA1 plays a key role in mediating and coordinating systemic cell-to-cell ROS and calcium signals that are required for plant acclimation to stress.

## INTRODUCTION

Reactive oxygen species (ROS; *i.e.,* H_2_O_2_, O_2_^.−^, ^1^O_2_, and HO^.^) are credited with playing a fundamental role in the evolution of life on Earth impacting processes such as the endosymbiotic event, emergence of multicellularity, and the development of reproduction through sex (Taverne et al., 2018; Hörandl and Speijer, 2018; Gutteridge and Halliwell, 2018; Jabłońska and Tawfik, 2021). Although originally considered to be toxic byproducts of aerobic metabolism, in recent years numerous studies revealed that ROS, such as H_2_O_2_ and O_2_^.−^, are essential for life, acting as key regulators of redox, stress responses, and cell-to-cell signaling (Schieber and Chandel, 2014; Mittler, 2017; Sies and Jones, 2020; Mittler et al., 2022). Examples for the role of ROS in cell-to-cell signaling include the recruitment of macrophages to wound sites and interactions between neurons in animals, communication between microorganisms within a microbiome, and transmission of long-distance cell-to-cell signals in plants (Aguirre and Lambeth, 2010; Razzell et al., 2013; Zheng et al., 2015; Zandalinas et al., 2020a; Zandalinas et al., 2020b; Fichman et al., 2021; Iwashita et al., 2021). In the flowering plant *Arabidopsis thaliana* (Arabidopsis), cell-to-cell signaling plays a pivotal role in local and systemic responses, acclimation, and survival of plants during stress (Mittler et al., 2011; Zhu, 2016; Waszczak et al., 2018; Smirnoff and Arnaud, 2019; Zandalinas et al., 2020a; Zandalinas et al., 2020b; Fichman et al., 2021; Mittler et al., 2022). During this process, ROS production is triggered in cells directly subjected to stress (termed ‘local tissue’), and a state of ‘activated ROS production’, driven by the function of respiratory burst oxidase homologs (RBOHs; the plant equivalents of mammalian NADPH oxidases; NOXs), is propagated from cell-to-cell over long distances, sometime spanning the entire length of the plant (Mittler et al., 2011; Zhu, 2016; Fichman et al., 2019; Zandalinas et al., 2020a; Fichman and Mittler, 2020b; Zandalinas et al., 2020b; Fichman et al., 2021; Mittler et al., 2022). Once the activated ROS production state reaches cells and tissues other than the ones initiating it (*i.e.,* tissues not directly subjected to stress; termed ‘systemic tissues’), it activates in them different acclimation mechanisms and enhances the overall resilience of the plant to stress (termed ‘systemic acquired acclimation’; SAA; Zandalinas et al., 2020a; Zandalinas et al., 2020b; Fichman et al., 2021). Although RBOHs such as RBOHD and RBOHF were found to produce ROS essential for this process (Miller et al., 2009; Fichman et al., 2019; Zandalinas et al., 2020a; Fichman and Mittler, 2020b; Zandalinas et al., 2020b; Fichman et al., 2021; Mittler et al., 2022), the identity of the ROS receptor(s) perceiving the ROS signal and enabling the cell-to-cell signaling process to occur is currently unknow.

We recently developed a new method for whole plant live ROS imaging to visualize cell-to-cell ROS signaling (termed the ‘ROS wave’), in mature plants growing in soil (Fichman et al., 2019; Fichman and Mittler, 2020b). Using this method, we screened over 120 different mutants, potentially involved in ROS and calcium signaling, for the presence or absence of the ROS wave in response to a local treatment of high light (HL) stress (table S1). Among the different mutants we screened were several putative receptors, including different cysteine-rich receptor-like kinases (CRKs) and the leucine-rich-repeat receptor-like kinase (LRR-RLK) HPCA1 (‘H_2_O_2_-induced Ca^2+^ increases 1’; At5g49760; also known as ‘cannot respond to DMBQ 1’; CARD1). HPCA1 was recently identified as a potential sensor for extracellular H_2_O_2_ (Wu et al., 2020). However, HPCA1 was also shown to function as a plant quinone receptor (Laohavisit et al., 2020). Here we reveal that HPCA1 acts as a key ROS receptor required for the propagation of cell-to-cell ROS signals, systemic signaling in response to different biotic and abiotic stresses, and plant acclimation to stress. We further show that HPCA1 is required for systemic calcium signals (also termed the ‘calcium wave’), but not systemic membrane depolarization responses (a type of ‘electric wave’), and that systemic calcium signals mediated by HPCA1 require the function of the calcium-permeable channel mechanosensitive ion channel like 3 (MSL3). In addition, we reveal that key components of calcium-dependent signaling cascades, such as the calcineurin B-like calcium sensor (CBL4), the CBL4-interacting protein kinase 26 (CIPK26), and the sucrose-non-fermenting-1-related protein kinase 2.6 (SnRK2.6, also termed ‘open stomata 1’, OST1), are also involved in this process. Our findings reveal that HPCA1 plays a key role in the sensing of H_2_O_2_ produced at the apoplast during cell-to-cell signaling, linking the accumulation of apoplastic H_2_O_2_ with calcium cascades and the activation of further ROS production by RBOHs; thereby mediating and coordinating systemic cell-to-cell ROS and calcium signals that are required for plant resilience to stress.

## RESULTS

### HPCA1 is required for systemic cell-to-cell ROS and calcium signaling during plant responses to HL stress

To study the role of HPCA1 in systemic cell-to-cell ROS signaling, we subjected a single leaf of wild-type (WT) and two independent knockout alleles of HPCA1 (*hpca1-1, hpca1-2*) to a HL stress treatment of 1700 µmol photons s^−1^m^−2^ for 2 min and used our newly developed whole-plant live ROS imaging method with 2’,7’-dichlorodihydrofluorescein diacetate (H_2_DCFDA) as a probe (Fichman et al., 2019) to measure the accumulation of ROS in local and systemic leaves over a period of 30 min. As shown in Fig. 1A, mutants deficient in HPCA1 (*hpca1-1, hpca1-2*) did not accumulate ROS in response to a local application of HL stress (see also movie S1). Because H_2_DCFDA detects a broad range of different ROS, we also used Peroxy Orange 1 (PO1; Fichman et al., 2019) instead of H_2_DCFDA as a probe, to measure the levels of H_2_O_2_ that accumulate in local and systemic leaves of WT, *hpca1-1,* and *hpca1-2* plants following a similar HL treatment. As shown in Fig. 1B, H_2_O_2_ accumulated in local and systemic leaves of WT, but not the *hpca1-1* and *hpca1-*2 mutants in response to a local treatment of HL stress.

**Fig. 1.**
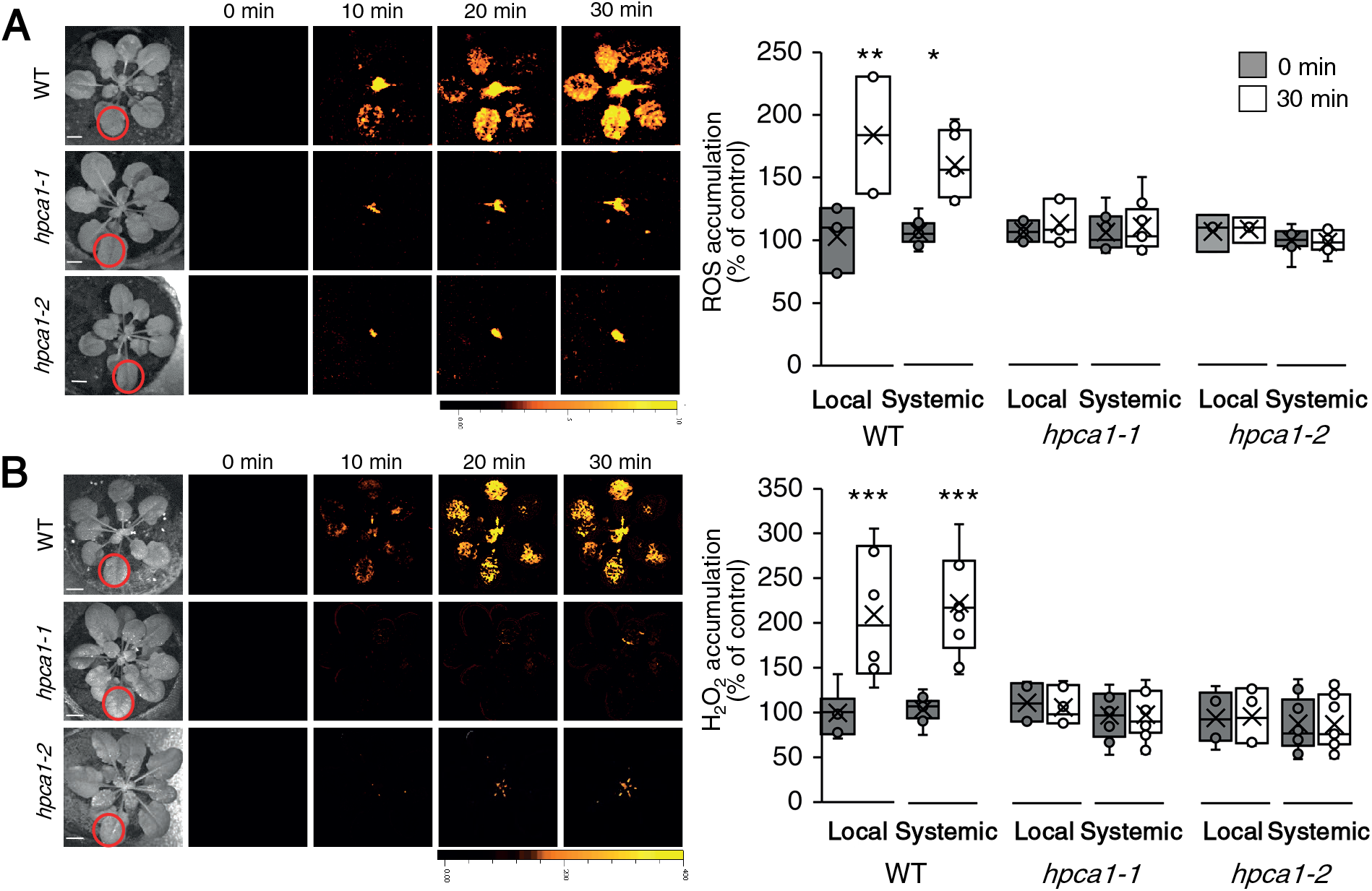
HPCA1 is required for systemic cell-to-cell ROS signaling in response to light stress. **(A)** *Arabidopsis* plants were subjected to a high light (HL) stress treatment applied to a single leaf (Local; indicated with a red circle), and ROS accumulation was imaged, using H_2_DCFDA, in whole plants (local and systemic tissues). Representative time-lapse images of whole plant ROS accumulation in WT, *hpca1-1* and *hpca1-2* plants are shown alongside bar graphs of combined data from all plants used for the analysis at the 0- and 30-min time points (local and systemic). **(B)** Same as in (A), but for whole plant H_2_O_2_ accumulation that was imaged using Peroxy Orange 1 (PO1). All experiments were repeated at least 3 times with 10 plants of each genotype per experiment. Data is presented as box plot graphs, where X is mean ± S.E., N=30, *P < 0.05, **P < 0.01, ***P < 0.001, Student t-test. Scale bar, 1 cm. See movie S1 for live imaging. Abbreviations: H_2_DCFDA, 2’,7’-dichlorodihydrofluorescein diacetate; HPCA1, H_2_O_2_-induced Ca^2+^ increases 1; PO1, Peroxy Orange 1; ROS, reactive oxygen species; WT, wild-type.

Upon sensing of H_2_O_2_, HPCA1 was found to trigger the accumulation of calcium in the cytosol (Wu et al., 2020). This process could activate another type of cell-to-cell signaling pathway termed the ‘calcium wave’ (dependent on the function of the calcium channels glutamate-like receptor 3.3 and 3.6; GLRs; Evans et al., 2016; Toyota et al., 2018; Shao et al., 2020; Fichman and Mittler, 2021a). To determine whether HPCA1 is also required for systemic cell-to-cell calcium signals, we subjected a single leaf of WT, *hpca1-1,* and *hpca1-2* plants to the same HL stress treatment described above and used Fluo-4-AM as a probe in our live imaging platform (Fichman and Mittler, 2021a) to measure changes in cytosolic calcium levels in local and systemic leaves over a period of 30 min. As shown in Fig. 2A, mutants deficient in HPCA1 (*hpca1-1, hpca1-2*) did not display local or systemic changes in cytosolic calcium levels in response to a local application of HL stress (see also movie S1).

**Fig. 2.**
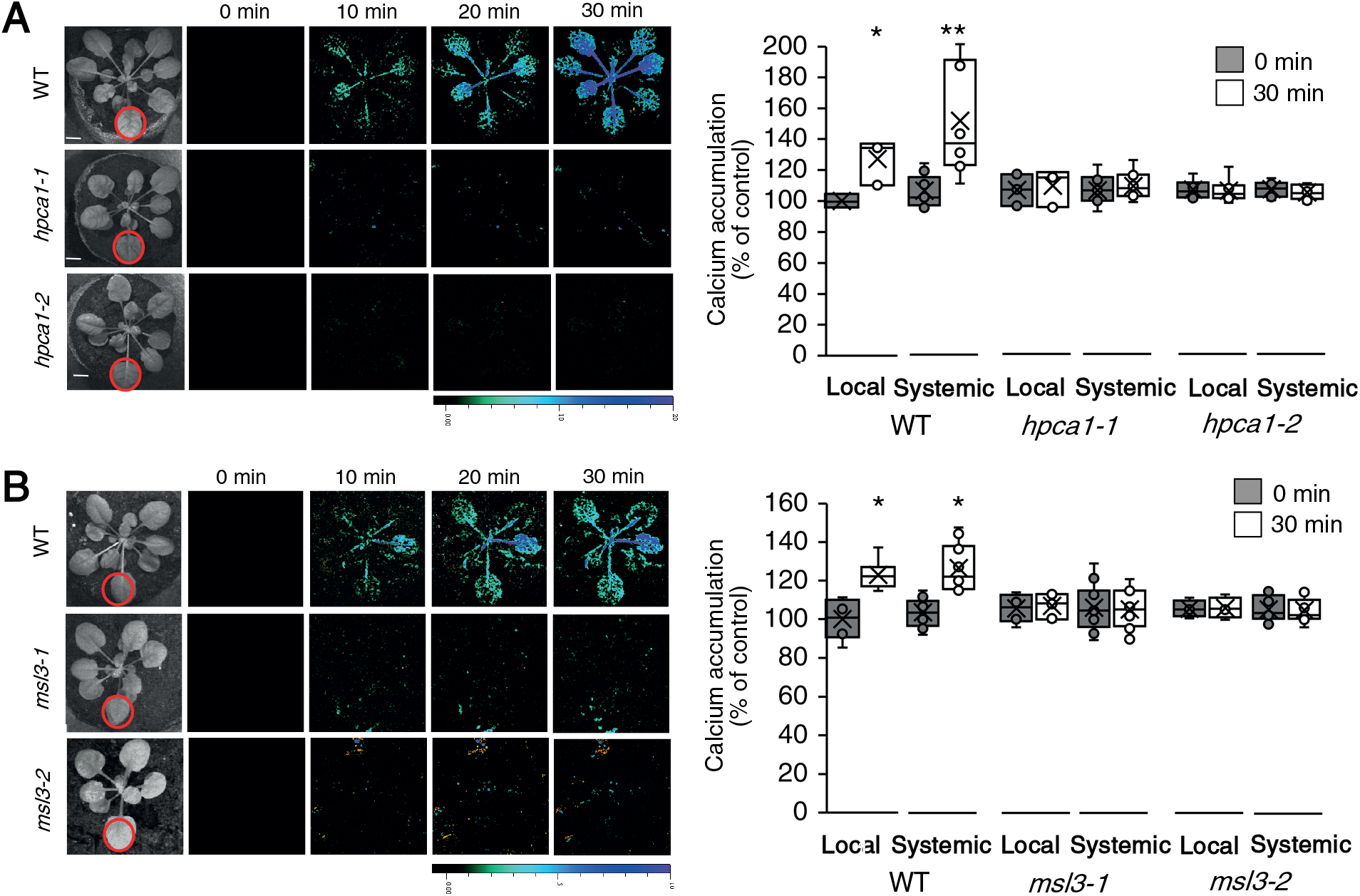
HPCA1 and MSL3 are required for systemic cell-to-cell calcium signaling in response to light stress. **(A)** *Arabidopsis* plants were subjected to a high light (HL) stress treatment applied to a single leaf (Local; indicated with a red circle), and cytosolic calcium accumulation was imaged using Fluo-4-AM in whole plants (local and systemic tissues). Representative time-lapse images of whole plant cytosolic calcium accumulation in WT, *hpca1-1* and *hpca1-2* plants are shown alongside bar graphs of combined data from all plants used for the analysis at the 0- and 30-min time points (local and systemic). **(B)** Same as in (A), but for WT, *msl3-1* and *msl3-2* plants. All experiments were repeated at least 3 times with 10 plants of each genotype per experiment. Data is presented as box plot graphs, where X is mean ± S.E., N=30, *P < 0.05, **P < 0.01, Student t-test. Scale bar, 1 cm. See movie S1 for live imaging. Abbreviations: HPCA1, H_2_O_2_-induced Ca^2+^ increases 1; MSL3, mechanosensitive ion channel like 3; WT, wild-type.

Systemic cell-to-cell ROS signals were previously found to be dependent on several different calcium-permeable channels including MSL3 (table S1; Fichman et al., 2021). We therefore used the method described above (Fig. 2A) to test whether systemic cell-to-cell cytosolic calcium changes are dependent on MSL3. As shown in Fig. 2B, in response to a local HL treatment, *msl3-1,* and *msl3-2* mutants did not display local or systemic changes in cytosolic calcium levels. This finding suggested that MSL3 could function downstream to HPCA1.

Systemic cell-to-cell calcium and ROS signals were previously proposed to be linked with another type of cell-to-cell signaling, termed the ‘electric wave’ (a rapid depolarization of the plasma membrane, also dependent on the function of GLRs; Mousavi et al., 2013; Nguyen et al., 2018; Farmer et al., 2020; Fichman and Mittler, 2021a). To determine whether HPCA1 is also required for systemic cell-to-cell membrane depolarization signals, we subjected a single leaf of WT, *hpca1-1,* and *hpca1-2* plants to the same HL stress treatment described above and used DiBAC_4_(3) as a probe in our live imaging platform (Fichman and Mittler, 2021a) to measure these changes in local and systemic leaves over a period of 30 min. Interestingly, while the systemic cell-to-cell calcium and ROS signals were suppressed in the *hpca1* mutants (Figs. 1, 2; movie S1), the rapid local and systemic membrane depolarization signal was not (Fig. 3; movie S1).

**Fig. 3.**
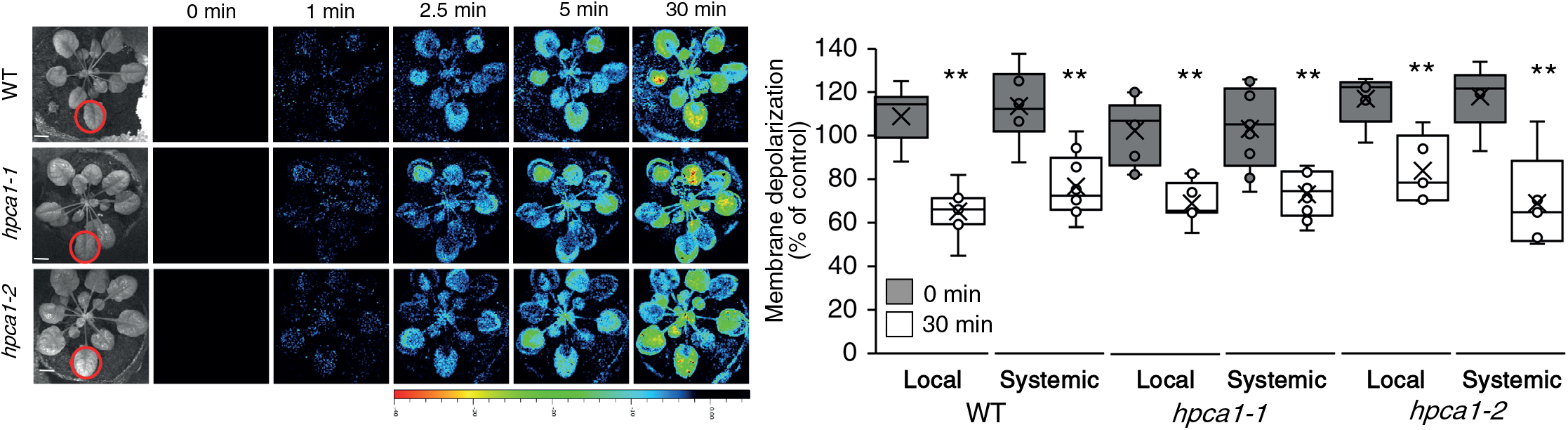
HPCA1 is not required for systemic cell-to-cell changes in membrane potential in response to light stress. *Arabidopsis* plants were subjected to a high light (HL) stress treatment applied to a single leaf (Local; indicated with a red circle), and changes in membrane potential were imaged using DiBAC_4_(3) in whole plants (local and systemic tissues). Representative time-lapse images of whole plant changes in membrane potential in WT, *hpca1-1* and *hpca1-2* plants are shown alongside bar graphs of combined data from all plants used for the analysis at the 0- and 30-min time points (local and systemic). All experiments were repeated at least 3 times with 10 plants of each genotype per experiment. Data is presented as box plot graphs where X is mean ± S.E., N=30, **P < 0.01, Student t-test. Scale bar, 1 cm. See movie S1 for live imaging. Abbreviations: DiBAC_4_(3), Bis-(1,3-Dibutylbarbituric Acid)Trimethine Oxonol; HPCA1, H_2_O_2_- induced Ca^2+^ increases 1; WT, wild-type.

### HPCA1 is required for local and systemic expression of different acclimation transcripts as well as for local and systemic plant acclimation to HL stress

Suppression of systemic cell-to-cell ROS and/or calcium signals (Figs. 1, 2) could prevent plants from acclimating to stress. To test whether HPCA1 mutants are deficient in plant acclimation, we measured the local and systemic expression of several transcripts associated with plant acclimation to excess light stress 30 min following the application of HL (1700 µmol photons s^−1^m^−2^) stress for 2 min to a local leaf of WT and *hpca1-1* plants. As shown in Fig. 4A, the expression of *MYELOBLASTOSIS DOMAIN PROTEIN 30 (MYB30), ZINC FINGER OF ARABIDOPSIS THALIANA 10 and 12 (ZAT10 and ZAT12)*, *ASCORBATE PEROXIDASE 2 (APX2)*, and *ZINC FINGER HOMEODOMAIN* 5 (ZHD5), was upregulated in local and systemic leaves of WT plants in response to the local HL stress treatment. In contrast, except for APX2 that was upregulated in local leaves of *hpca1-1* plants, the expression of all transcripts was suppressed in local and systemic leaves of *hpca1-1* plants in response the local HL stress treatment (Fig. 4A).

**Fig. 4.**
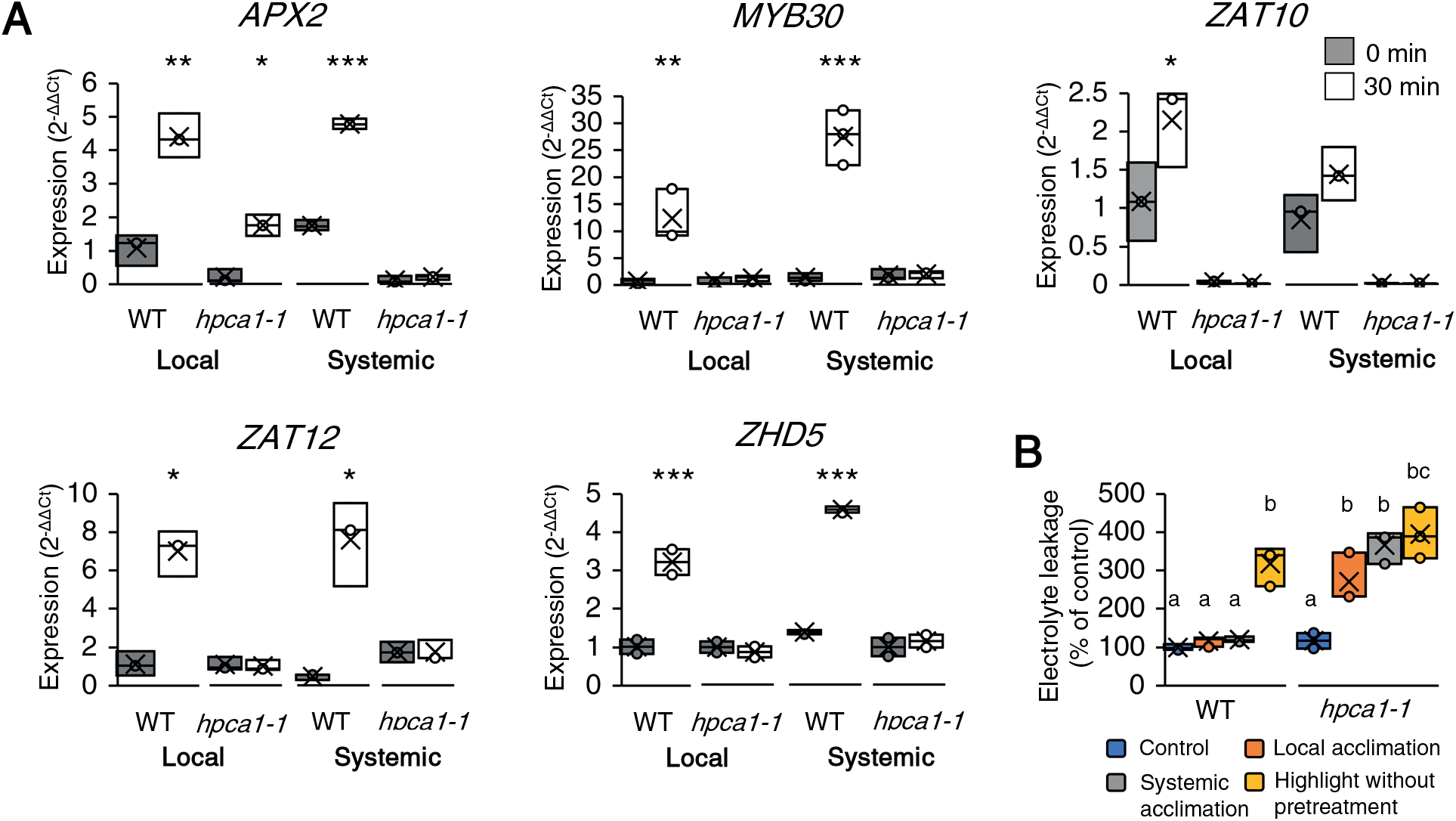
HPCA1 is required for local and systemic expression of stress-acclimation transcripts, as well as acclimation of plants to light stress. **(A)** Real-time quantitative PCR analysis of *APX2*, *MYB30*, *ZAT10*, *ZAT12*, and *ZHD5* expression in local and systemic leaves of wild-type and *hpca1-1* plants subjected to a local HL treatment. Transcripts tested were previously found to respond to HL stress in wild-type plants. Results are presented as relative quantity (RQ) compared to control WT from local leaf. **(B)** Averaged measurements of leaf injury (increase in ion leakage) of WT and *hpca1-1* plants. Measurements are shown for unstressed plants (control), local leaves subjected to a pretreatment of HL stress before a long HL stress period (local acclimation), systemic leaves of plants subjected to a local HL stress pretreatment before a long period of local HL stress was applied to a systemic leaf (systemic acclimation), and systemic leaves of plants subjected to a long HL stress period without pretreatment (HL without pretreatment). Results are presented as percent of control (leaves not exposed to HL stress). All experiments were repeated at least 3 times with 10 plants of each genotype per experiment. Data is presented in (A) as box plot graphs where X is mean ± S.E., N=30, *P < 0.05, **P < 0.01, ***P < 0.001, Student t-test. Data is presented in (B) as box plot graphs where X is mean ± S.E, N=30, one-way ANOVA followed by a Tukey test; lowercase letters donate significance (p < 0.05). Abbreviations: *APX2*, *ASCORBATE PEROXIDASE 2*; HL, high light; HPCA1, H_2_O_2_-induced Ca^2+^ increases 1; *MYB30*, *MYELOBLASTOSIS DOMAIN PROTEIN 30*; PCR, polymerase chain reaction; WT, wild-type; *ZAT10*, *ZINC FINGER OF ARABIDOPSIS THALIANA 10*; *ZAT12, ZINC FINGER OF ARABIDOPSIS THALIANA 12; ZHD5, ZINC FINGER HOMEODOMAIN* 5.

The lack of systemic ROS and calcium cell-to-cell signals (Figs. 1, 2), as well as systemic expression of *MYB30*, *ZAT10*, *ZAT12*, *APX2* and *ZHD5* (Fig. 4A), could suggest that HPCA1 is required for systemic acclimation of plants to HL stress. To test this possibility, we measured the acclimation (*i.e.,* reduced tissue damage following exposure to light stress) of mature WT and *hpca1-1* plants to a prolonged HL stress treatment following a short pretreatment with HL stress and an incubation period. As shown in Fig. 4B, pretreatment of WT plants with 10 min of HL stress, followed by an incubation of 50 min under controlled growth conditions, protected plants from a subsequent exposure to 45 min of HL stress (*i.e.,* prevented leaf injury as measured by electrolyte leakage, compared to plants that were subjected to the 45 min HL treatment without a 10 min pretreatment with excess white or red light). In contrast, pretreatment of *hpca1-1* plants with a short HL stress failed to induce plant acclimation to a subsequent prolonged HL stress that resulted in a significant increase in electrolyte leakage from cells (Fig. 4B).

### HPCA1 is required for the propagation but not initiation of systemic cell-to-cell ROS signal

Systemic cell-to-cell ROS signaling is driven by two different pathways, one that controls its initiation at the local tissue, and one that controls its propagation-, amplification-, and acclimation-promoting functions, in local and systemic tissues (Fichman et al., 2021; Mittler et al., 2022). These two pathways can be distinguished in plants by grafting experiments between WT plants and different mutants (Fichman et al., 2021). Using such grafting experiments, we found that HPCA1 is required for the propagation but not initiation of the ROS wave (Fig. 5). Thus, while the *hpca1-1* mutant was deficient in ROS wave propagation through the scion (systemic tissue), following the activation of the ROS wave at the WT stock (local tissue), it could transmit the systemic signal through the (local) stock tissue to a WT scion (Fig. 5A-5C). In contrast, the *rbohD* mutant was deficient in both systemic cell-to-cell ROS signal initiation and propagation (Fig. 5D; (Fichman et al., 2021), while the *rbohF* mutant was similar to *hpca1-1* mutant and was only deficient in ROS wave propagation (Fig. 5E). HPCA1 is therefore required for the propagation of the systemic cell-to-cell ROS signal (Fig. 5A-5C), as well as for its transcript accumulation- and acclimation-driven functions in systemic tissues (Fig. 4).

**Fig. 5.**
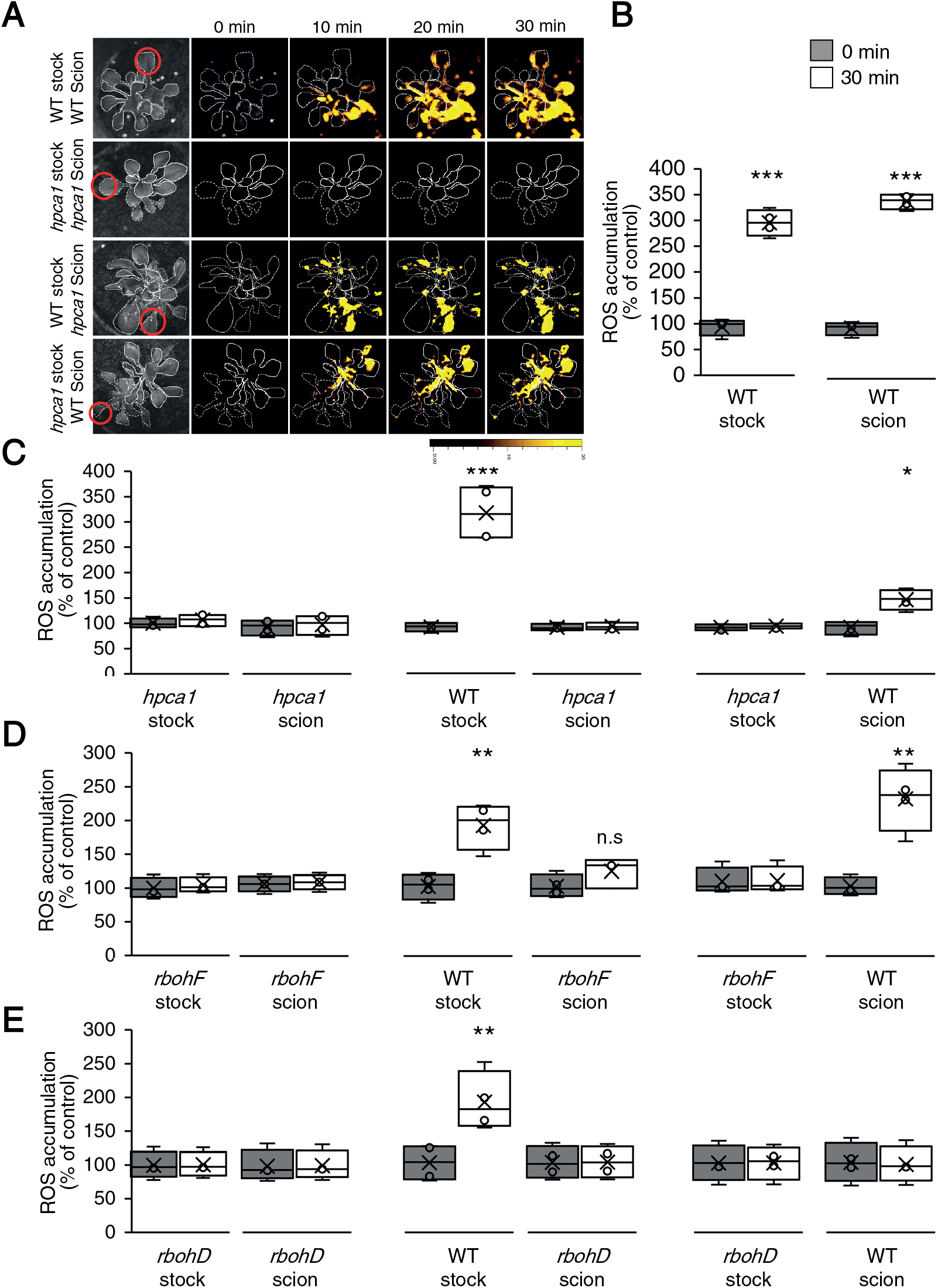
HPCA1 is required for systemic cell-to-cell ROS signal propagation, but not initiation, in response to light stress. **(A)** Representative time-lapse images of ROS accumulation in stock and scion parts of grafted plants, generated using WT and *hpca1-1* plants, in response to HL stress applied to a single leaf (indicated with a red circle) belonging to the stock part. Scions are indicated by solid white lines, and stocks are indicated by dashed white lines. **(B)** Bar graphs showing the combined data from the stock and scion of grafted WT plants subjected to HL stress on a single leaf of the stock scion. **(C)** Same as (B), but for different grafting combinations between WT and *hpca1-1* plants. **(D)** Same as (B), but for different grafting combinations between WT and *rbohF* plants. **(E)** Same as (B), but for different grafting combinations between WT and *rbohD* plants. All experiments were repeated at least 3 times with 10 plants of each genotype per experiment. Data is presented as box plot graphs where X is mean ± S.E., N=30, *p < 0.05, **P < 0.01, ***P < 0.001, Student t-test. Abbreviations: HL, high light; HPCA1, H_2_O_2_-induced Ca^2+^ increases 1; *rbohD,* respiratory burst oxidase homolog D; *rbohF,* respiratory burst oxidase homolog F; ROS, reactive oxygen species; WT, wild-type.

### HPCA1 is required for systemic cell-to-cell ROS signaling in response to biotic and abiotic stresses, but not in response to wounding

The findings that HPCA1 is required for the propagation of the ROS wave (Fig. 5A-5C), that plays a key role in plant responses to many different abiotic stresses (Zhu, 2016; Fichman et al., 2019; Zandalinas et al., 2020a; Fichman and Mittler, 2020b; Zandalinas et al., 2020b; Fichman et al., 2021; Mittler et al., 2022), could suggest that HPCA1 is involved in plant responses to a broad range of stresses. To test the involvement of HPCA1 in local and systemic ROS responses to other stresses, we treated a local leaf of WT or *hpca1-1* plants with a bacterial pathogen (Fichman et al., 2019), salt stress, or wounding (Fichman et al., 2019), and measured local and systemic accumulation of ROS (untreated, or mock buffer treatment in the absence of the pathogen or salt were used as controls). As shown in Fig. 6, while all treatments caused the accumulation of ROS in local and systemic leaves of WT plants, *hpca1-1* plants did not respond to the bacterial pathogen or salt stress treatments (Fig. 6A, 6B). In response to a local treatment of wounding, *hpca1-1* mutants did however display a local and systemic cell-to-cell ROS signaling response that was indistinguishable from that of WT (Fig. 6C). These findings suggest that cell-to-cell ROS signals could be mediated in plants by more than one type of ROS receptor. Systemic cell-to-cell ROS signaling pathways, triggered by HL stress, bacterial infection, or salinity treatments (Figs. 1, 6A, 6B) and mediated by HPCA1, could therefore be distinguished from those activated by wounding (Fig. 6C) and potentially mediated by a yet unknown ROS receptor(s).

**Fig. 6.**
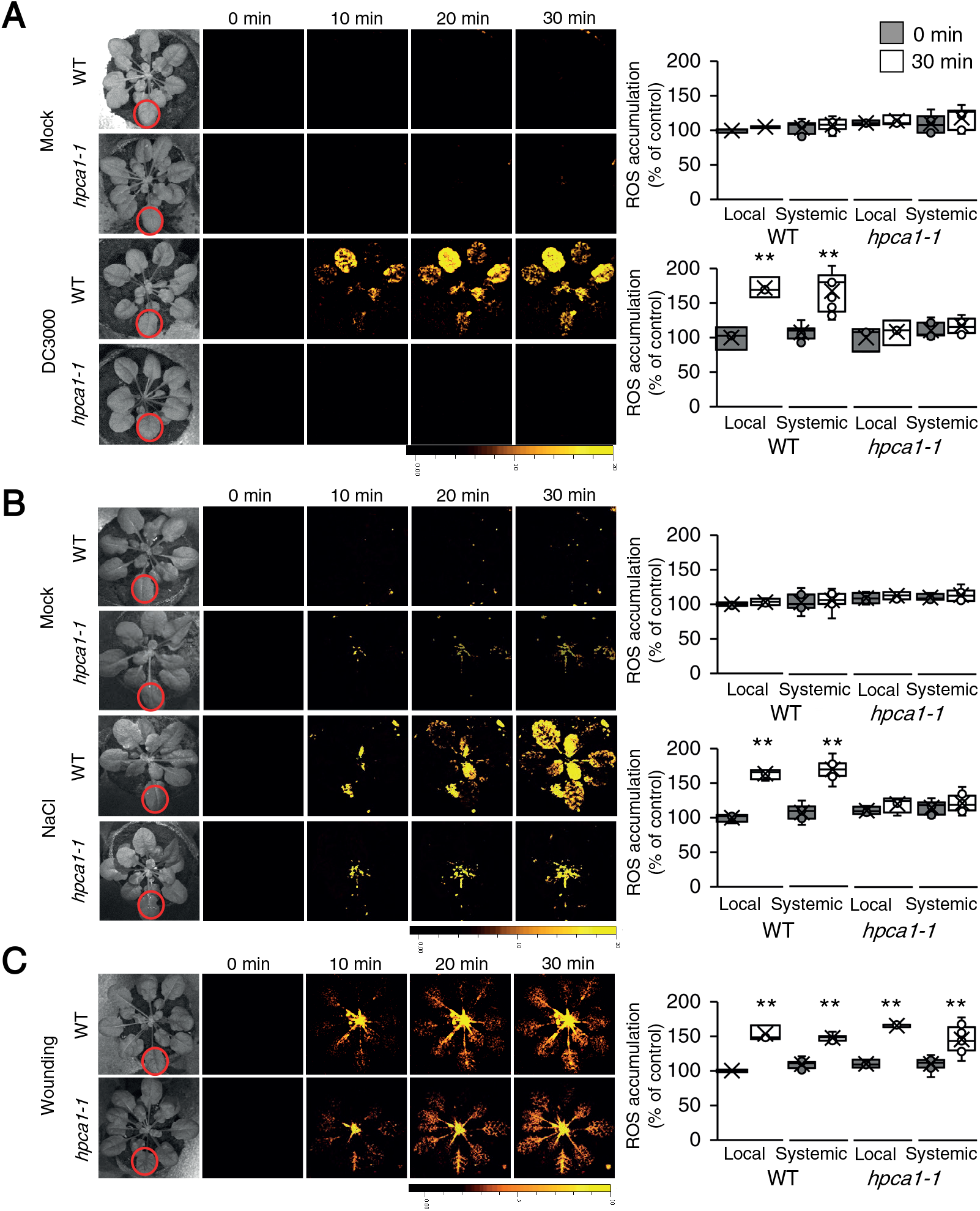
HPCA1 is required for systemic cell-to-cell ROS responses to bacterial infection and salt stress, but not wounding. **(A)** Representative time-lapse images of whole plant ROS accumulation in WT and *hpca1-1* plants subjected to mock or bacterial (*Pseudomonas syringae* DC3000) infection on a single local leaf are shown alongside bar graphs of combined data from all plants used for the analysis at the 0- and 30-min time points (local and systemic). **(B)** Same as in (A), but for mock and salt stress (100 mM NaCl) applied to a single local leaf. **(C)** Same as in (A), but for wounding applied to a single local leaf (control plants were untreated). All experiments were repeated at least 3 times with 10 plants of each genotype per experiment. Data is presented as box plot graphs where X is mean ± S.E., N=30, **P < 0.01, Student t-test. Abbreviations: HPCA1, H_2_O_2_-induced Ca^2+^ increases 1; ROS, reactive oxygen species; WT, wild-type.

### HPCA1-dependent cell-to-cell ROS signaling requires the central calcium signaling regulators CBL4, CIPK26, and OST1

The increase in calcium levels resulting from HPCA1 activation during local and systemic responses to HL stress (Fig. 2) could cause the activation of calcium-dependent protein kinase cascades and trigger ROS production by RBOHs (Luan and Wang, 2021; Mittler et al., 2022). Our mutant screen (table S1) identified three proteins potentially involved in such cascades (CBL4, CIPK26, and OST1). As shown in Fig. 7A, similar to the *hpca1-1* mutant (Fig. 1), *cbl4-1*, *cipk26-2*, and *ost1-2* mutants were deficient in mediating the systemic cell-to-cell ROS signal in response to a 2 min local treatment of HL stress. In addition, and also similar to the *hpca1-1* mutant (Fig. 4B), *cbl4-1*, *cipk26-2*, and *ost1-2* mutants were unable to acclimate to HL stress following a pretreatment with a short period of HL stress (Fig. 7B).

**Fig. 7.**
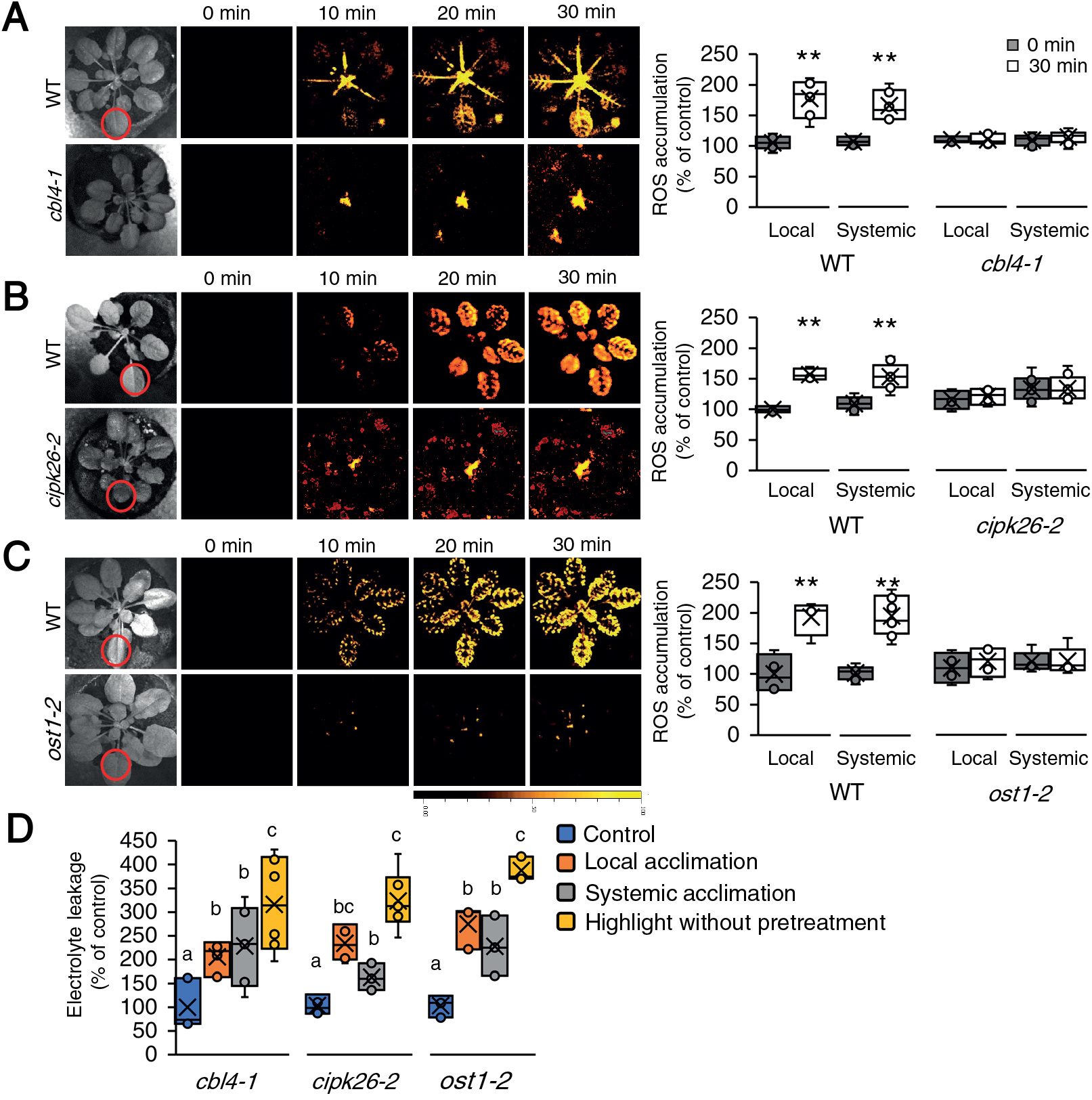
CBL4, CIPK26, and OST1 are required for systemic cell-to-cell ROS signaling and acclimation to light stress. **(A)** Representative time-lapse images of whole plant ROS accumulation in wild-type (WT) and *cbl4-1* plants subjected to a local HL stress treatment (applied to a single local leaf; indicated with a red circle) are shown alongside bar graphs of combined data from all plants used for the analysis at the 0- and 30-min time points (local and systemic). **(B)** Same as (A), but for WT and *cipk26-2* plants. **(C)** Same as (A), but for WT and *ost1-2* plants. **(D)** Averaged measurements of leaf injury (increase in ion leakage) in WT, *cbl4*, *cipk26*, and *ost1* plants. Measurements are shown for unstressed plants (control), local leaves subjected to a pretreatment of HL stress before a long HL stress period (local acclimation), systemic leaves of plants subjected to a local HL stress pretreatment before a long period of local HL stress was applied to a systemic leaf (systemic acclimation), and systemic leaves of plants subjected to a long HL stress period without pretreatment (HL without pretreatment). All experiments were repeated at least 3 times with 10 plants of each genotype per experiment. Data is presented in (A) to (C) as box plot graphs where X is mean ± S.E., N=30, **P < 0.01, Student t-test. Data is presented in (D) as box plot graphs where X is mean ± S.E., N=30, one-way ANOVA followed by a Tukey test; lowercase letters donate significance (p < 0.05). Abbreviations: CBL4, calcineurin B-like calcium sensor 4; CIPK26, CBL4-interacting protein kinase 26; HL, high light; OST1, open stomata 1; ROS, reactive oxygen species; WT, wild-type.

To test whether CBL4, CIPK26, and OST1 are required for the initiation or propagation of the systemic cell-to-cell ROS signal, we conducted grafting experiments between these mutants and WT plants (Fig. 8; similar to the analysis described in Fig. 5). These studies revealed that similar to HPCA1 (Fig. 5), CBL4, CIPK26, and OST1 are all required for the propagation but not initiation of the systemic cell-to-cell ROS signal. Thus, while the *cbl4-1*, *cipk26-2*, and *ost1-2* mutants were deficient in ROS wave propagation through the scion (systemic tissue), following the activation of the ROS wave at the WT stock (local tissue), they could transmit the systemic signal through the (local) stock tissue to a WT scion (Fig. 8).

**Fig. 8.**
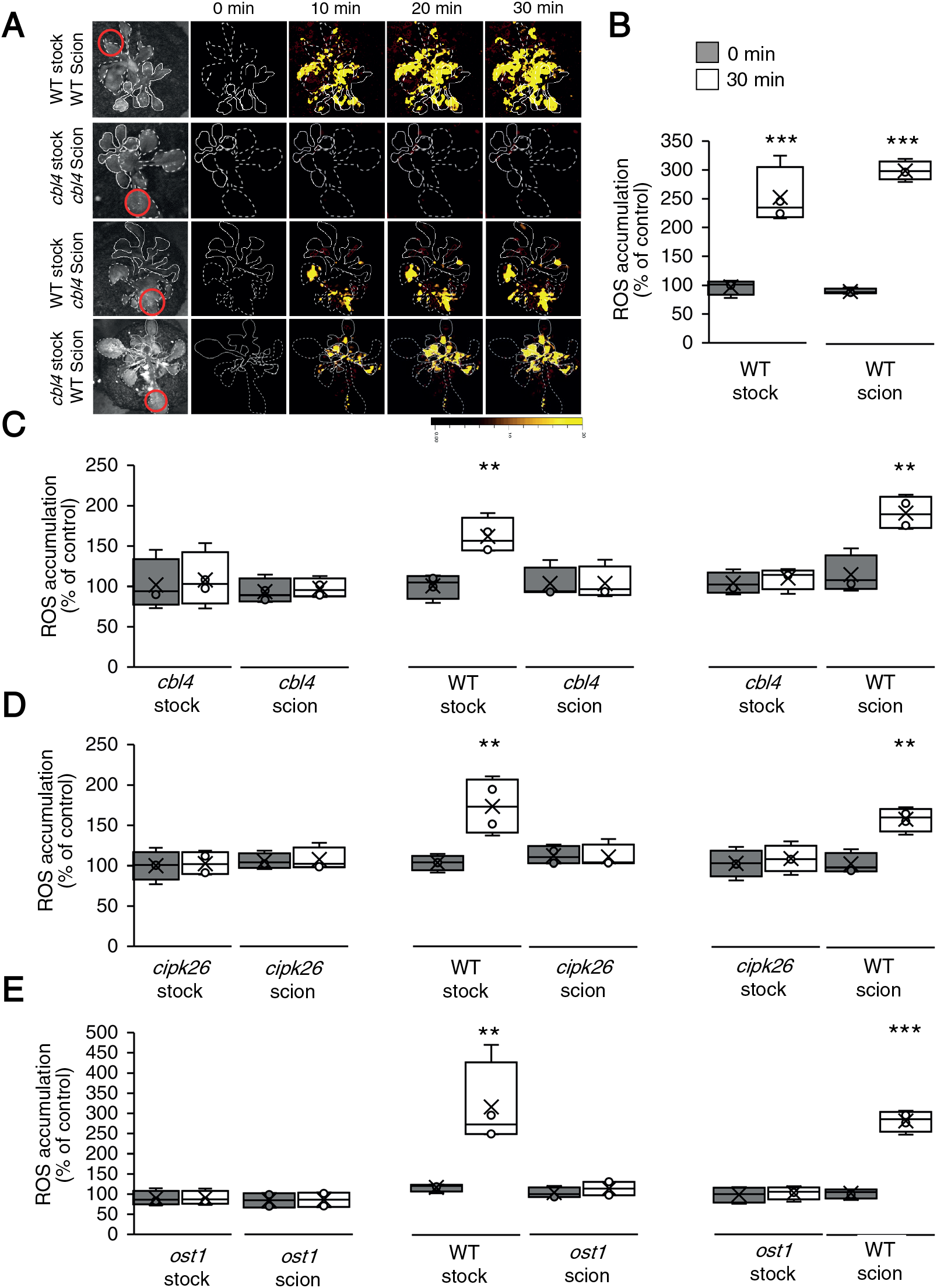
CBL4, CIPK26, and OST1 are required for systemic ROS signal propagation, but not initiation, in response to light stress. **(A)** Representative time-lapse images of ROS accumulation in stock and scion parts of grafted plants, generated using WT and *cbl4-1* plants, in response to a local HL stress treatment applied to a single leaf (indicated with a red circle) belonging to the stock part. Scions are indicated by solid white lines, and stocks are indicated by dashed white lines. **(B)** Bar graphs showing the combined data from the stock and scion of grafted WT plants subjected to HL stress on a single leaf of the stock scion. **(C)** Same as (B), but for different grafting combinations between WT and *cbl4-1* plants. **(D)** Same as (B), but for different grafting combinations between WT and *cipk26-2* plants. **(E)** Same as (B), but for different grafting combinations between WT and *ost1-2* plants. All experiments were repeated at least 3 times with 10 plants of each genotype per experiment. Data is presented as box plot graphs where X is mean ± S.E., N=30, **P < 0.01, ***P < 0.001, Student t-test. Abbreviations: CBL4, calcineurin B-like calcium sensor 4; CIPK26, CBL4-interacting protein kinase 26; HL, high light; OST1, open stomata 1; ROS, reactive oxygen species; WT, wild-type.

The findings that key components of a calcium-dependent signaling cascade (*i.e.,* CBL4, CIPK26, and OST1) are required for the propagation of the cell-to-cell ROS signal reveal that enhanced levels of calcium alone (Fig. 2) are not sufficient to trigger the ROS wave by directly interacting with the calcium-binding domains of RBOHD (Ogasawara et al., 2008). Rather, an amplification cascade of the signal is needed.

### The same amino acid residue required for RBOHD activation by OST1 is also required for RBOHD activation during systemic cell-to-cell ROS signaling

The sensing of high cytosolic calcium levels by CBL4 was shown to activate CIPK26, and CIPK26 was shown to phosphorylate and activate RBOHF and OST1 (Drerup et al., 2013). OST1, in turn, was shown to phosphorylate RBOHD on serine 347 and activate it (Wang et al., 2020). OST1 was also shown to phosphorylate and activate RBOHF (Sirichandra et al., 2009). Because RBOHD plays such a canonical role in the initiation and propagation of the systemic cell-to-cell ROS signal (Fig. 5; Zandalinas et al., 2020a; Zandalinas et al., 2020b; Fichman et al., 2021), we tested whether deleting its N-terminal regulatory domain (RD; amino acids 1 to 347), or mutating serine 347 to alanine (the target of OST1 phosphorylation; Wang et al., 2020), will inhibit the systemic cell-to-cell ROS signal in response to HL stress. For this purpose, we expressed the WT *RbohD* gene (RbohD genomic; Fig. 9), or the *RbohD* cDNA (RbohD cDNA; Fig. 9), under the control of the *RbohD* promoter in *rbohD* mutants. In addition, we expressed the *RbohD* cDNA without the RD (RbohD w/o RD; Fig. 9), or the RbohD gene with point mutations (Serine to Alanine) in positions 22 and 26 (RbohD S22-26A; Fig. 9), or 22, 26, 343 and 347 (RbohD S22-26,343-347A; Fig. 9) in the *rbohD* mutant (Nühse et al., 2007; Zandalinas et al., 2020b). Phosphorylation of RBOHD on S343/S347, as well as on S22/S26 was previously associated with the RBOHD- and ROS-dependent innate immune response of Arabidopsis (with S343/S347 playing a key role in this response; Nühse et al., 2007), and the WT *RbohD* gene expressed under the control of the *RbohD* promoter was shown to complement local and systemic ROS production in response to HL stress in the *rbohD* mutant (Zandalinas et al., 2020b). Once we confirmed that all transgenic complementation assays were homozygous and expressing a single copy of the transgene, we subjected a single leaf of WT, *rbohD*, *rbohD*RbohD genomic, *rbohD*RbohD cDNA, *rbohD*RbohD w/o RD, *rbohD*RbohD S22-26A, and *rbohD*RbohD S22-26,343-347A to a 2 min of HL stress treatment (as described for Fig. 1) and measured ROS accumulation in local and systemic leaves. As shown in Fig. 9A and 9B, complementation of the *rbohD* mutant with the WT *RbohD*, WT *RbohD* cDNA, or *RbohD* S22-26A restored the systemic cell-to-cell ROS response. In contrast, complementation of the *rbohD* mutant with the *RbohD* w/o RD, or the *RbohD* S22-26,343-347A failed to restore the systemic ROS signal.

**Fig. 9.**
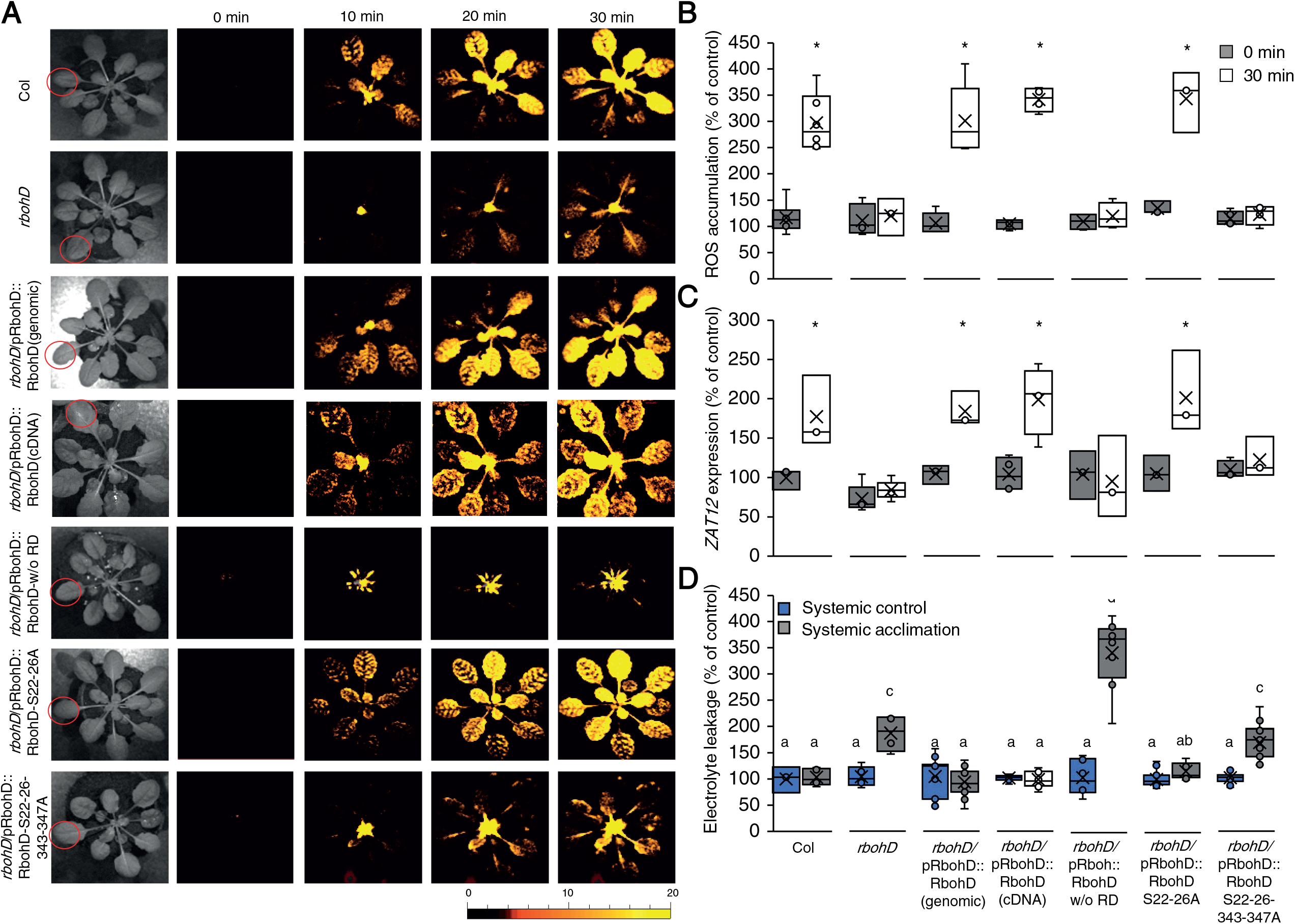
Mutating specific amino acids in RBOHD suppresses systemic ROS accumulation in response to high light stress. **(A)** Representative time-lapse images of whole plant ROS accumulation in WT, *rbohD*, *rbohD* complemented with the wild type *RbohD* gene [*rbohD*/pRbohD::RbohD (genomic)], *rbohD* complemented with the *RbohD* cDNA expressed under the control of the *RbohD* promoter [*rbohD*/pRbohD::RbohD (cDNA)], *rbohD* complemented with the *RbohD* cDNA without the N-terminal regulatory domain (RD, 1-347) expressed under the control of the *RbohD* promoter [*rbohD*/pRboh::RbohD w/o RD], rbohD complemented with the *RbohD* gene with S22A and S26A mutations [*rbohD*/pRbohD::RbohD S22-26A], or *rbohD* complemented with the *RbohD* gene with S22A, S26A, S343A and S347A mutations [*rbohD*/pRbohD::RbohD S22-26-343-347A], following treatment of a single local leaf with HL stress (indicated with a red circle). **(B)** Bar graphs of combined data from all plants used for the analysis shown in (A) at the 0- and 30-min time points (systemic). **(C)** Bar graphs of combined *Zat12* promoter activity (luciferase imaging) in systemic leaves of *rbohD*/*Zat12::luciferase* double homozygous plants transformed with all vectors shown in (A), measured at 0- and 30-min time following application of HL stress to a single local leaf. **(D)** Averaged measurements of leaf injury (increase in ion leakage) in systemic tissues of all lines shown in (A). Measurements are shown for unstressed systemic leaves (systemic control) and systemic leaves of plants subjected to a local HL stress pretreatment before a long period of local HL stress was applied to a systemic leaf (systemic acclimation). All experiments were repeated at least 3 times with 10 plants of each genotype per experiment. Two independent transgenic lines for each construct were averaged. Data presented in (B) and (C) is mean ± S.E., N=30, *P < 0.05, Student t-test. Data presented in (D) is mean ± S.E., N=30, one-way ANOVA followed by a Tukey test; lowercase letters donate significance (p < 0.05). Abbreviations: cDNA, complementary DNA; HL, high light; *RbohD,* respiratory burst oxidase homolog D; RD, regulatory domain; ROS, reactive oxygen species; WT, wild-type; *Zat12*, Zinc finger of *Arabidopsis thaliana* 12.

To study the expression of the key HL acclimation response gene *Zat12* in *rbohD* mutants transformed with the different constructs, we conducted the same analysis described above, however instead of the *rbohD* mutant we used the double homozygous line expressing the *Zat12::luciferase* reporter in the *rbohD* background (developed as described in Miller et al., 2009; Zandalinas et al., 2020b) for the complementation study. As shown in Fig. 9C, expression of the *Zat12* gene (measured by luciferase activity; Miller et al., 2009; Zandalinas et al., 2020b) was significantly elevated only in *rbohDZat12::luciferase* lines complemented with the WT *RbohD*, WT *RbohD* cDNA, or *RbohD* S22-26A (as well as in WT plants transformed with the *Zat12::luciferase* reporter). In contrast, *Zat12* expression was not complemented in *rbohDZat12::luciferase* lines by expression of the *RbohD* w/o RD or the *RbohD* S22-26,343-347A constructs. These findings agreed with the measurements of local and systemic ROS shown for the different complemented *rbohD* lines in panels A and B.

To study systemic acclimation to HL stress we also subjected the *rbohD* complemented lines (Fig. 9A, 9B) to the same HL SAA assay shown in Figs. 4B and 7B. As shown in Fig. 9D, complementation of the *rbohD* mutant with the WT *RbohD*, WT *RbohD* cDNA, or *RbohD* S22-26A restored systemic HL acclimation to the *rbohD* mutant, while complementation of the *rbohD* mutant with the *RbohD* w/o RD or the *RbohD* S22-26,343-347A construct did not.

Taken together, the analyses shown in Fig 9 suggest that complementation of *rbohD* with the wild type *RbohD* gene, cDNA, or *RbohD* gene with mutations in S22 and S26 (Nühse et al., 2007; Zandalinas et al., 2020b), restored HL-induced systemic cell-to-cell ROS signaling (Fig. 4A), systemic *Zat12* gene expression (Fig. 4B), and systemic acclimation to HL stress (Fig. 4C). By contrast, complementation of *rbohD* with the *RbohD* cDNA that lacks the RD, or the *RbohD* gene that contains point mutations in S22, S26, S343 and S347 (Nühse et al., 2007), did not restore the ROS wave, systemic *Zat12* expression, or systemic acclimation to HL (Fig. 9). These findings point to residues S343 and S347 (the target of OST1; Wang et al., 2020) as playing a key role in cell-to-cell ROS signaling.

## DISCUSSION

The ability of plants to mobilize a signal from a small group of cells subjected to stress to the entire plant, *i.e.,* systemic signaling, plays a pivotal role in plant acclimation to, and/or defense against, many different abiotic and biotic stresses (Mittler et al., 2011; Zhu, 2016; Waszczak et al., 2018; Smirnoff and Arnaud, 2019; Farmer et al., 2020; Johns et al., 2021; Mittler et al., 2022). Among the different signal transduction mechanisms that mediate systemic responses in plants is a rapid cell-to-cell signaling process that involves membrane depolarization, cytosolic calcium alterations, and ROS accumulation (Figs. 1-3, movie S1; Mittler et al., 2011; Farmer et al., 2020; Fichman and Mittler, 2020a; Shao et al., 2020; Johns et al., 2021; Mittler et al., 2022). Previous studies identified RBOHD, RBOHF, and GLR3.3GLR3.6 as key players in this cell-to-cell response (Miller et al., 2009; Mousavi et al., 2013; Toyota et al., 2018; Shao et al., 2020; Zandalinas et al., 2020b). While RBOHs were shown to mediate ROS production required for cell-to-cell signaling and plant acclimation (Miller et al., 2009; Fichman et al., 2019), GLRs were shown to mediate membrane depolarization and alterations in calcium levels (that could potentially drive ROS production; Mousavi et al., 2013; Evans et al., 2016; Toyota et al., 2018; Nguyen et al., 2018; Shao et al., 2020; Fichman and Mittler, 2021a). Prior studies have also suggested that the function of RBOHs and GLRs is interlinked (*e.g.,* Fichman and Mittler, 2020a; Fichman and Mittler, 2021a). Nevertheless, how changes in ROS levels at the apoplast (produced by RBOHs) are translated into changes in calcium in the cytosol of neighboring cells remains unknown. Here we show that HPCA1 plays a canonical role in systemic cell-to-cell signaling in plants, triggering cytosolic calcium accumulation upon sensing of apoplastic ROS/H_2_O_2_ (Figs. 1, 2A, movie S1). The altered calcium levels, potentially driven by MSL3 (Fig. 2B), could then activate a downstream pathway that requires CBL4, CIPK26, and OST1 and trigger further ROS production (Figs. 7-9). HPCA1 may therefore represent a highly important and missing puzzle piece that links changes in apoplastic ROS levels driven by RBOH function with changes in cytosolic calcium levels driven by different calcium-permeable channels such as MSL3 (Fig. 10). The finding that HPCA1 is required for systemic ROS and calcium cell-to-cell signaling (Figs. 1, 2A), the expression of many acclimation transcripts in systemic tissues (Fig. 4A), as well as plant acclimation (Fig. 4B), provides strong support to this proposed role of HPCA1.

**Fig. 10.**
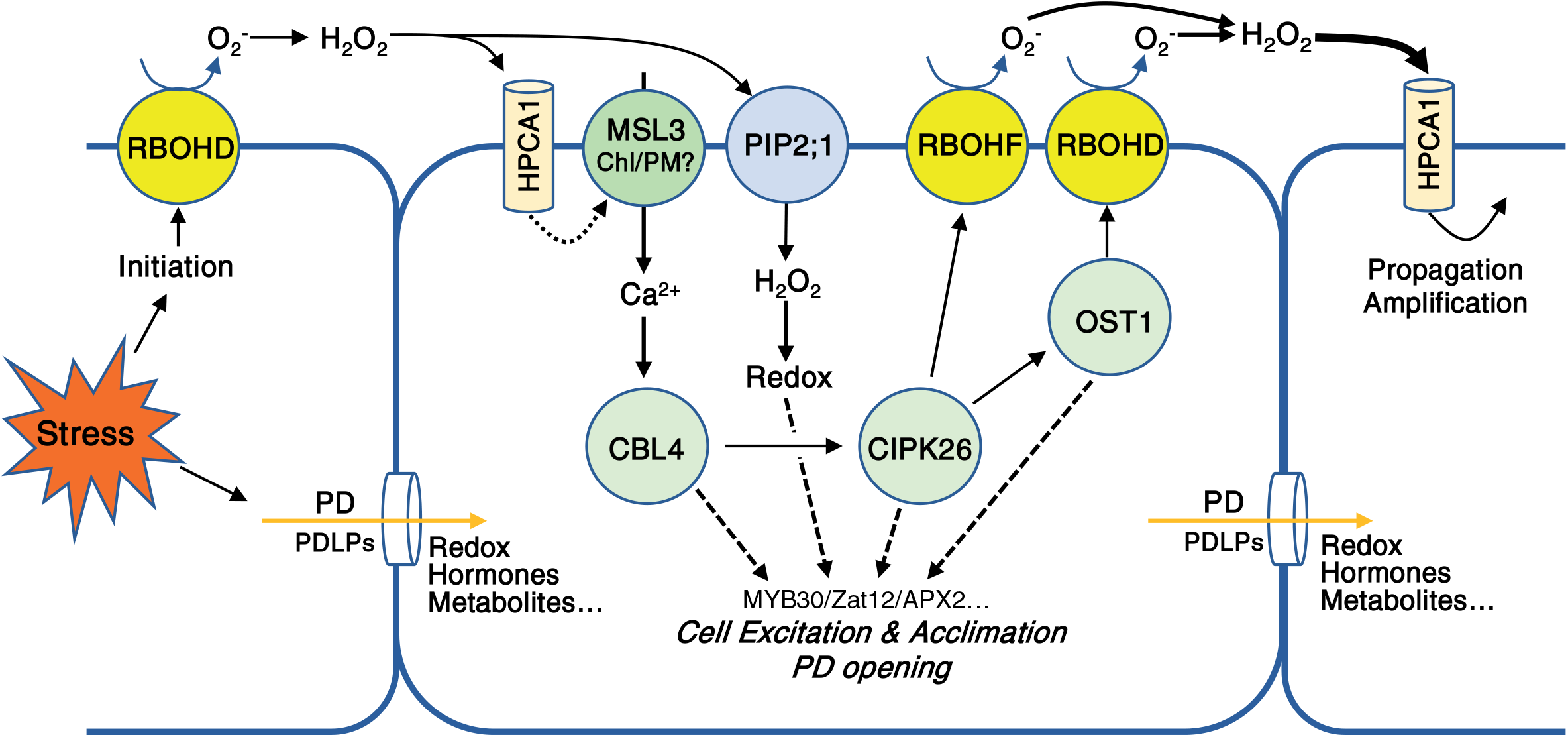
A model depicting the role of HPCA1 in the propagation of cell-to-cell ROS signaling in plants. HPCA1 is proposed to sense ROS at the apoplast and trigger an increase in cytosolic calcium levels via MSL3. The increase in calcium is proposed to activate a kinase cascade involving CBL4, CIPK26 and OST1 that together activate RBOHD and RBOHF enhancing ROS production at the apoplast; that is sensed by the HPCA1 of the next cell in the cell-to-cell chain of ROS signaling. ROS that accumulate in the apoplast (mainly H_2_O_2_) are shown to enter the cell via aquaporins and alter the redox state of different transcriptional regulators. The function of the pathway activated by HPCA1 is shown to be required for the enhanced transcript expression, acclimation, and resilience of plants to stress (please see text for more details). Abbreviations: APX2, Ascorbate peroxidase 2; HPCA1, H_2_O_2_-induced Ca^2+^ increases 1; CBL4, calcineurin B-like calcium sensor 4; CIPK26, CBL4-interacting protein kinase 26; MYB30, Myeloblastosis domain protein *30*; OST1, open stomata 1; PD, plasmodesmata; PDLP, plasmodesmata localized protein; RBOHD, respiratory burst oxidase homolog D, RBOHF, respiratory burst oxidase homolog F; ROS, reactive oxygen species; ZAT12, Zinc finger of *Arabidopsis thaliana* 12

Interestingly, in our hands, HPCA1 appears not to be needed for the mediation of systemic membrane potential changes (Fig. 3; movie S1). In this respect it should also be noted that our grafting experiments (Fig. 5) revealed that the mobilization of the systemic cell-to-cell signal through a scion made from *hpca1-1* (without the accumulation of detectable ROS levels) could lead to the activation of the ROS cell-to-cell signal in the WT scion (Fig. 5). Taken together, these findings suggest that a cell-to-cell membrane potential signals could mediate a systemic signal even in the absence of the ROS and/or calcium cell-to-cell signals. This notion is also supported by the pace of the different systemic signals detected in our study (Figs 1-3; movie S1). The systemic change in membrane potential (a type of electric wave) is the fastest, followed by a change in cell-to-cell cytosolic calcium levels, that are followed by changes in cell-to-cell ROS levels (Figs. 1-3; movie S1). These observations could suggest that an electric wave (that is GLR-dependent, at least for its initiation; Mousavi et al., 2013; Nguyen et al., 2018; Fichman and Mittler, 2021a) is the first to reach all cells. The changes in membrane potential it brings with it may prime, alter, or activate different channels and other signaling mechanisms. These could then trigger a calcium wave [that could be dependent on GLRs, MSLs, two-pore channel 1 (TPC1), and/or cyclic nucleotide–gated ion channels (CNGCs); Evans et al., 2016; Toyota et al., 2018; Shao et al., 2020; Fichman et al., 2021; Dickinson et al., 2022)], that in turn activate ROS production via CBL4-, CIPK26- and/or OST1-mediated RBOH activation (Figs 7-10). Although calcium changes are imaged in our system before ROS changes (Figs. 1-3; movie S1), the new player in this pathway, introduced by this work, *i.e.,* HPCA1, appears to be required for integrating the cell-to-cell calcium and ROS signals, providing a mechanistic understanding to how changes in apoplastic ROS levels are linked to changes in cytosolic calcium levels (Fig. 10; movie S1). The possible role of electric signals in activating cell-to-cell ROS signaling is also supported by a recent study showing that aboveground plant-to-plant transmission of electric signals (via two physically touching leaves) can trigger the cell-to-cell ROS signal in a receiving plant, and that this communication process is dependent on GLRs, RBOHs and MSLs (Szechynska-Hebda et al., 2022).

Interestingly, although HPCA1 was found to be required for systemic cell-to-cell ROS responses to local HL, salt, or pathogen treatments (Figs. 1, 6), it was not required for cell-to-cell ROS signaling in response to wounding (Fig. 6). This finding suggested that different receptors for apoplastic ROS could be involved in mediating systemic cell-to-cell signaling in response to different stresses. Alternatively, the sensing of changes in apoplastic ROS levels may not play a key role in systemic cell-to-cell signaling in response to wounding. In this respect it should be noted that in addition to being sensed at the plasma membrane by HPCA1, ROS (H_2_O_2_) can also enter the cytosol from the apoplast through aquaporins (Rodrigues et al., 2017; Fichman et al., 2021; Fig. 10). A recent study has shown for example that in the aquaporin mutant plasma membrane intrinsic protein 2;1 (*pip2;1*), the cell-to-cell ROS signal triggered by HL stress is suppressed (Fichman et al., 2021; Mittler et al., 2022). ROS could also move from cell-to-cell via plasmodesmata that open in an RBOHD-dependent manner during the progression of the cell-to-cell signal (Fichman et al., 2021). Further studies are needed to address the relationships between apoplastic sensing of ROS via HPCA1, cytosolic sensing of ROS following their entry into the cell via aquaporins, and the transfer of ROS from cell-to-cell via plasmodesmata (Fig. 10; Fichman et al., 2021).

An overall view of rapid cell-to-cell ROS and calcium signaling emerges from our study. In this view each cell in the cell-to-cell ROS signaling pathway senses the ROS generated by the cell preceding it via HPCA1, activates a calcium-dependent signal transduction pathway (involving CBL4, CIPK26 and OST1), and triggers ROS production by RBOHD and RBOHF (Fig. 10). The activation of ROS production by that cell is then sensed by the cell following it in the chain, via its own HPCA1, and the process is repeated forming a positive amplification loop that drives the ROS signal from cell-to-cell until all cells in the plant turn their ROS production state to ‘activated’. While the initiation of the cell-to-cell ROS signal is primarily dependent on RBOHD (Miller et al., 2009; Fichman et al., 2019), its propagation is dependent on RBOHD and RBOHF (Fig. 5), that together could amplify the ROS signal (Fig. 10). CIPK26 can activate RBOHF and OST1 (Drerup et al., 2013), while OST1 can activate RBOHD and RBOHF (Sirichandra et al., 2009; Wang et al., 2020; Figs. 7-10). Activation of HPCA1 could also cause the opening of aquaporins such as PIP2;1 (Rodrigues et al., 2017; Smirnoff and Arnaud, 2019; Maurel et al., 2021; Mittler et al., 2022) and facilitate the transfer of RBOH-generated ROS into cells. The enhanced production of apoplastic ROS by each cell could therefore alter the ROS and redox state of the cytosol (Fichman and Mittler, 2021b), in an aquaporin- and plasmodesmata-dependent manner (Fichman et al., 2021), and activate multiple transcriptional regulators such as MYB30 and ZAT12 (Fig. 4; Mittler et al., 2022), causing all cells ‘excited’ or ‘activated’ by the cell-to-cell ROS signal to acquire a heightened state of tolerance to the stress and become acclimated (Figs. 4, 7, 9, 10; Zandalinas et al., 2020a; Zandalinas et al., 2020b; Fichman et al., 2021; Fichman and Mittler, 2021b; Mittler et al., 2022). Cell-to-cell ROS signaling therefore plays a key role in plant acclimation to stress, and HPCA1 is a key component of this pathway enabling ROS sensing and continued signal propagation (Fig. 10).

## Materials and Methods

### Plant material, growth conditions and generation of transgenic plants

*Arabidopsis thaliana* Col-0 wild type plants, homozygous knockout lines (Alonso et al., 2003) of *hpca1* (AT5G49760; CS923304), *cbl4* (AT5G24270; CS859749; Yang et al., 2019), *cipk26* (AT5G21326; SALK_074944C; Lyzenga et al., 2013), *ost1* (AT4G33950; SALK_020604), *msl3* (SALK_201695C; CS69719), *rbohD* (AT5G47910; CS68747; Torres et al., 2002), and *rbohF* (AT1G64060; CS68748; Torres et al., 2002), as well as native promoter complementation lines of *rbohD* with full-length genomic sequence of *RBOHD*, *RBOHD* S22-26A, *RBOHD* S22-26-343-347A; Nühse et al., 2007), cDNA sequence of *RBOHD* (Zandalinas et al., 2020b) and cDNA sequence of *RBOHD* without its regulatory domain (ΔM1-S347; generated as described below) were used for the main figures (additional mutants are described in table S1). Plants were grown in peat pellets (Jiffy International, Kristiansand, Norway) under controlled conditions of 10hr/14hr light/dark regime, 50 µmol photons s^−1^ m^−2^ and 21°C for 4 weeks (Zandalinas et al., 2020a; Zandalinas et al., 2020b; Fichman et al., 2021). For constructing RBOHD without the regulatory domain (ΔM1- S347), a DNA fragment lacking the *RbohD* regulatory domain (from amino acid 348 to 921) was amplified by PCR from cDNA template (using specific primers: 5’-GAGACTCGAGATGCAGAAGCTTAGACCGGCAAA-3’ and 5’-TCTCGAGCTCCTAGAAGTTCTCTTTGTGGAAGT-3’), isolated and sequenced. The resulting *RbohD* sequence without its regulatory domain was cloned into pCAMBIA2301 vectors (Marker Gene Technologies, Eugene, OR, USA) downstream of the native *RbohD* promoter (Nühse et al., 2007; Zandalinas et al., 2020b) replacing the full-length cDNA sequence of *RbohD* (using XhoI and SacI). *Agrobacterium tumefaciens* GV3101 (Koncz and Schell, 1986) was transformed with the binary plasmid and transgenic Arabidopsis plants were generated using floral dipping (Clough and Bent, 1998). Transformed seedlings were selected on 0.5X Murashige and Skoog media plates (Caisson Labs, Smithfield, UT, USA) supplemented with 50 µg ml^−1^ Kanamycin (Gold Bio, St. Louis, MO, USA) for three generations. Transgenic double homozygous pZat12::Luc *rbohD* plants (Miller et al., 2009; Zandalinas et al., 2020b) were also complemented with the different *RbohD* constructs (*i.e.,* full-length genomic sequence of *RbohD*, *RbohD* S22-26A, *RbohD* S22-26-343-347A, cDNA sequence of *RbohD*, and *RbohD* ΔM1-S347) as described above.

### Grafting

Grafting was performed as previously described (Fichman et al., 2021). Briefly, Arabidopsis plants (wild-type and different mutants) were germinated on 0.5X Murashige and Skoog media plates (Caisson Labs, Smithfield, UT, USA). An incision was made in seven-day-old stock seedlings to insert a scion into the cut while keeping the rosette of the stock plant intact. Plants were grown for five days in growth chamber at 20°C under constant light. Surviving grafted plants were transplanted to peat pellets and grown as described above for 5 days before light stress treatment (applied to a single leaf of the stock). For each knockout line, four combinations were constructed and tested: wild-type (WT) as the scion and the stock, the mutant line as the scion and the stock, mutant scion on WT stock, and WT scion on a mutant stock. Grafting was repeated 40 times for each combination of each line with approximately 40% success rate.

### Stress application and imaging of ROS, calcium and membrane potential

As previously described (Fichman et al., 2019; Zandalinas et al., 2020b; Fichman and Mittler, 2021a; Fig. S1), plants were fumigated for 30 min with 50 µM H_2_DCFDA (Millipore-Sigma, St. Louis, MO, USA) for ROS imaging (Fichman et al., 2019; Zandalinas et al., 2020b), 4.5 µM Fluo-4-AM (Becton, Dickinson and Company, Franklin Lakes, NJ, USA) for calcium imaging (Fichman and Mittler, 2021a), 20 µM DiBAC_4_(3) (Biotium, Fermont, CA, USA) for membrane potential imaging (Fichman and Mittler, 2021a), or 100 µM Peroxy Orange 1 (PO1; Millipore-Sigma, St. Louis, MO, USA) for H_2_O_2_ imaging (Fichman et al., 2019), using a nebulizer (Punasi Direct, Hong Kong, China) in a glass container. Following fumigation, different stresses were applied as described in (Fichman et al., 2019; Zandalinas et al., 2020b; Fichman and Mittler, 2021a). Briefly, plants were subjected to HL stress by illuminating a single leaf with 1700 µmol photons s^−1^m^−2^ using a ColdVision fiber optic LED light source (Schott, Southbridge, MA, USA; Fichman et al., 2019; Zandalinas et al., 2020b); pathogen infection was performed by dipping a single leaf in a solution containing DCF and *P. syringae* DC 3000 or the same solution without the bacteria (mock; Fichman et al., 2019); for wounding, a single leaf was pierced simultaneously by 20 dresser pines (Fichman et al., 2019; Fichman and Mittler, 2021a); for salt stress, a single leaf was dipped in 100 mM NaCl, 50 mM phosphate buffer, pH 7.4, with 50 µM H_2_DCFDA for 5 seconds (the same solution without NaCl was used for mock control). Fluorescence images were acquired using IVIS Lumina S5 (PerkinElmer, Waltham, MA, USA) for 30 min. ROS, H_2_O_2_, and calcium accumulation, as well as membrane depolarization were analyzed using Living Image 4.7.2 software (PerkinElmer, Waltham, MA, USA) using the math tools (Fichman et al., 2019; Zandalinas et al., 2020b; Fichman and Mittler, 2021a). Time course images were generated and radiant efficiency of regions of interest (ROI) were calculated. Each data set includes standard error of 8-12 technical repeats.

### Systemic acquired acclimation and electrolyte leakage assays

Local and systemic acquired acclimation to HL stress were measured by subjecting a local leaf to light stress (1700 µmol photons s^−1^m^−2^) for 0 or 10 min, incubating the plant under controlled conditions for 50 min, and then exposing the same leaf (local) or a younger leaf (systemic) to HL stress (1700 µmol photons s^−1^m^−2^) for 45 min (Zandalinas et al., 2020b; Fichman et al., 2021). Electrolyte leakage was measured by immersing the sampled (treated, untreated, local, or systemic) leaf in distilled water for 1 hr and measuring the conductivity of the water using Oakton CON 700 conductivity meter (Thermo Fisher Scientific, Vernon Hills, IL, USA). Samples were then boiled with the water, cooled down to room temperature and measured again for conductivity (total leakage). Electrolyte leakage was calculated as percentage of the conductivity before heating the samples over that of the boiled samples and compared between plants treated for 10 min on local leaf (pretreated) or treated for 0 min on their local leaf (non-pretreated). Experiments consisted of 5 repeats for each condition in each line. Standard error was calculated using Microsoft Excel; one-way ANOVA (confidence interval = 0.05) and Tukey honestly significant difference (HSD) were performed with IBM SPSS 25.

### Transcript expression

Transcript expression in response to HL stress in local and systemic leaves was measured using 4-week-old wild type and *hpca1-1* plants following the application of HL to a single leaf for 2 min (Fichman and Mittler, 2021a; Fichman et al., 2021). Exposed leaf (local) and unexposed fully developed younger leaf (systemic) were collected for RNA extraction at 0- and 30-min. RNA was extracted using Plant RNeasy kit (Qiagen, Hilden, Germany) according to the manufacture instructions. Total RNA was used for cDNA synthesis (PrimeScript RT Reagent Kit; Takara Bio, Takara Bio, Kusatsu, Japan). Transcript expression was quantified by real-time qPCR using iQ SYBR Green supermix (Bio-Rad Laboratories, Hercules, CA, USA), as previously described (Fichman and Mittler, 2021a; Fichman et al., 2021), with the following primers:s

*APX2* (AT3G09640) 5’-TCATCCTGGTAGACTGGACAAA-3’ and 5’-CACATCTCTTAGATGATCCACACC-3’; *MYB30* (AT3G28910) 5’- CCACTTGGCGAAAAAGGCTC-3’ and 5’- ACCCGCTAGCTGAGGAAGTA-3’; *ZAT10* (AT1G27730) 5’- ACTAGCCACGTTAGCAGTAGC-3’ and 5’- GTTGAAGTTTGACCGGAAGTC-3’; *ZAT12* (AT5G59820) 5’- TGGGAAGAGAGTGGCTTGTTT-3’ and 5’- TAAACTGTTCTTCCAAGCTCCA-3’; *ZHD5* (AT1G75240) 5’ - CCACCAATCCAAGTCTCCCTC-3’ and 5’-GCTCGCCGCATGATTCTTTAG-3’ and *Elongation factor 1 alpha* (5’-GAGCCCAAGTTTTTGAAGA-3’ and 5’-TAAACTGTTCTTCCAAGCTCCA-3’) was used for normalization of relative transcript levels. Results in the exponent of base 2 delta-delta terminal cycle were obtained by normalizing the relative transcript and comparing it to control WT from local leaf. Data represents 12 biological repeats and 3 technical repeats for each reaction. Standard error and Student t-test were calculated with Microsoft Excel.

### *ZAT12* promoter activity

Expression of luciferase driven by the *ZAT12* promoter was detected by luminescence imaging (Miller et al., 2009; Zandalinas et al., 2020b). Plants were sprayed with 1 mM luciferin (Gold Bio, St. Louis, MO, USA), and a single leaf was exposed to HL stress for 2 min (1700 µmol photons s^−1^m^−2^; ColdVision fiber optic LED light source; Schott, Southbridge, MA, USA). Plants were then imaged with the IVIS Lumina S5 apparatus (PerkinElmer, Waltham, MA, USA), as described before (Zandalinas et al., 2020b). Results are presented as precent of control (0 min). Each data set includes standard error of 8-12 technical repeats.

### Statistical analysis

All experiments were repeated at least three times with at least three biological repeats. Graphs were generated with Microsoft Excel and are box plots with x as mean ± SE. P values (*p < 0.05, **P < 0.01, ***P < 0.001) were generated with two-tailed Student t-test paired samples. ANOVA followed by a Tukey’s HSD post hoc test was used for hypothesis testing (different letters denote statistical significance at p < 0.05).

## Acknowledgements

We thank The Arabidopsis Biological Resource Center (ABRC), and Professors E.E. Farmer, S. Karpinski, E. Liscum, C. Maurel, S. Pandey, A. S. Richter, G. Stacey and S. Zhang for seeds that were used for the screens shown in table S1.

## Author contribution

Conceptualization: RM, YF, SIZ; Investigation: YF, SIZ; Visualization: YF, SIZ; Funding acquisition: RM; Resources: SL, SP, RM; Writing – original draft: RM, YF; Writing – review & editing: RM, YF, SIZ, SL, SP.

## Data and materials availability

All data and materials are available upon request from RM.

## Supplementary Materials

**Table S1.**
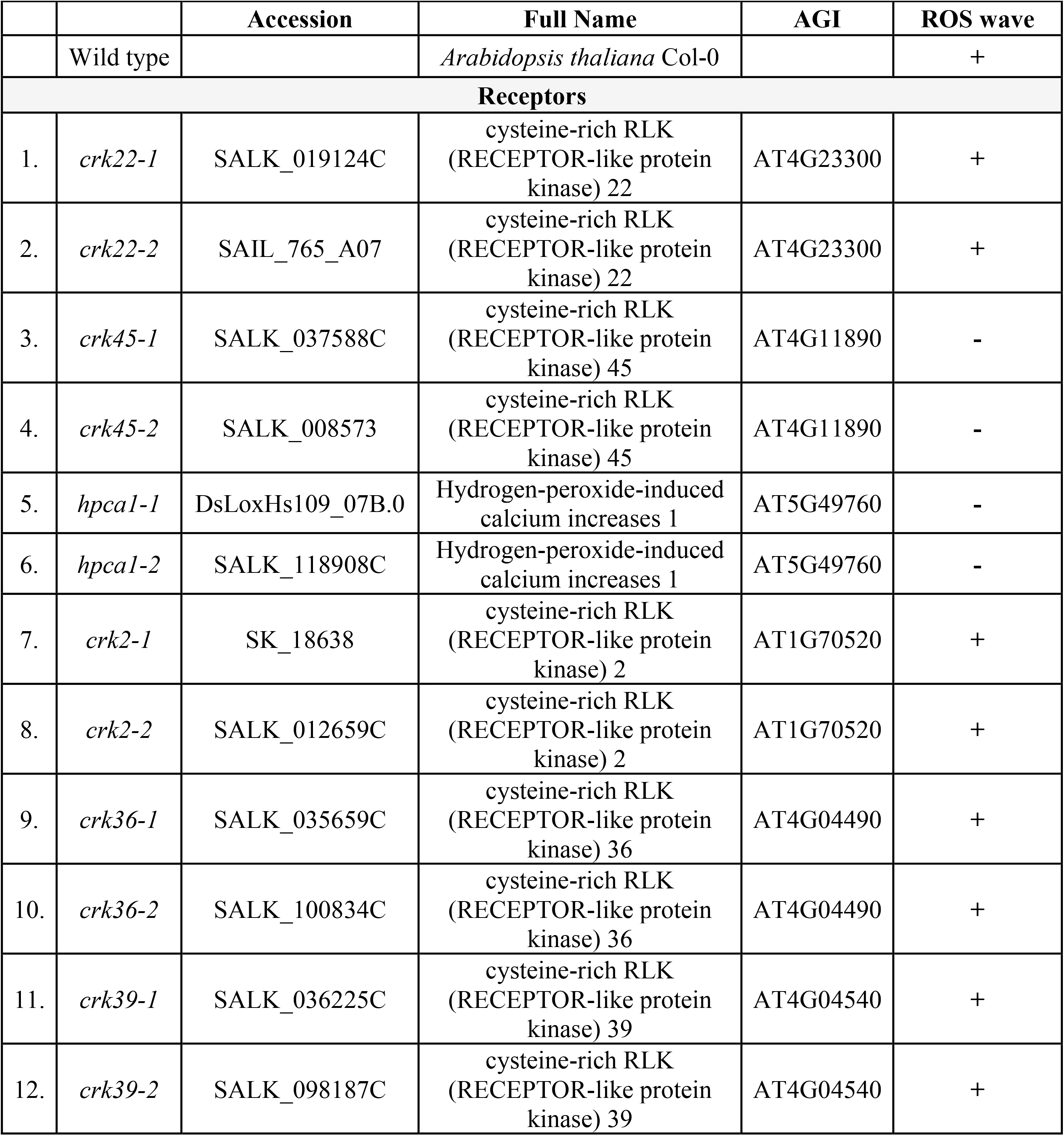
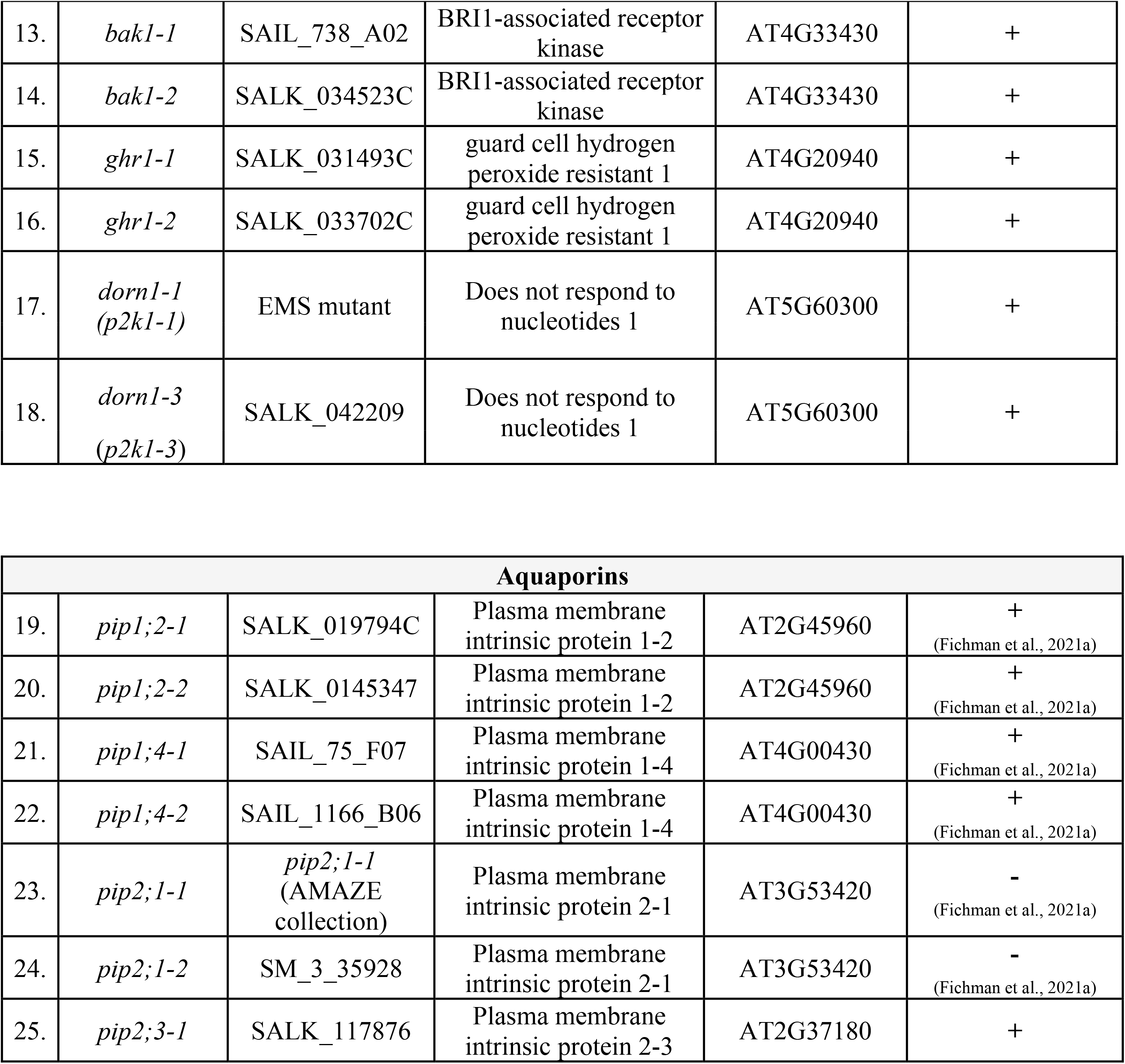
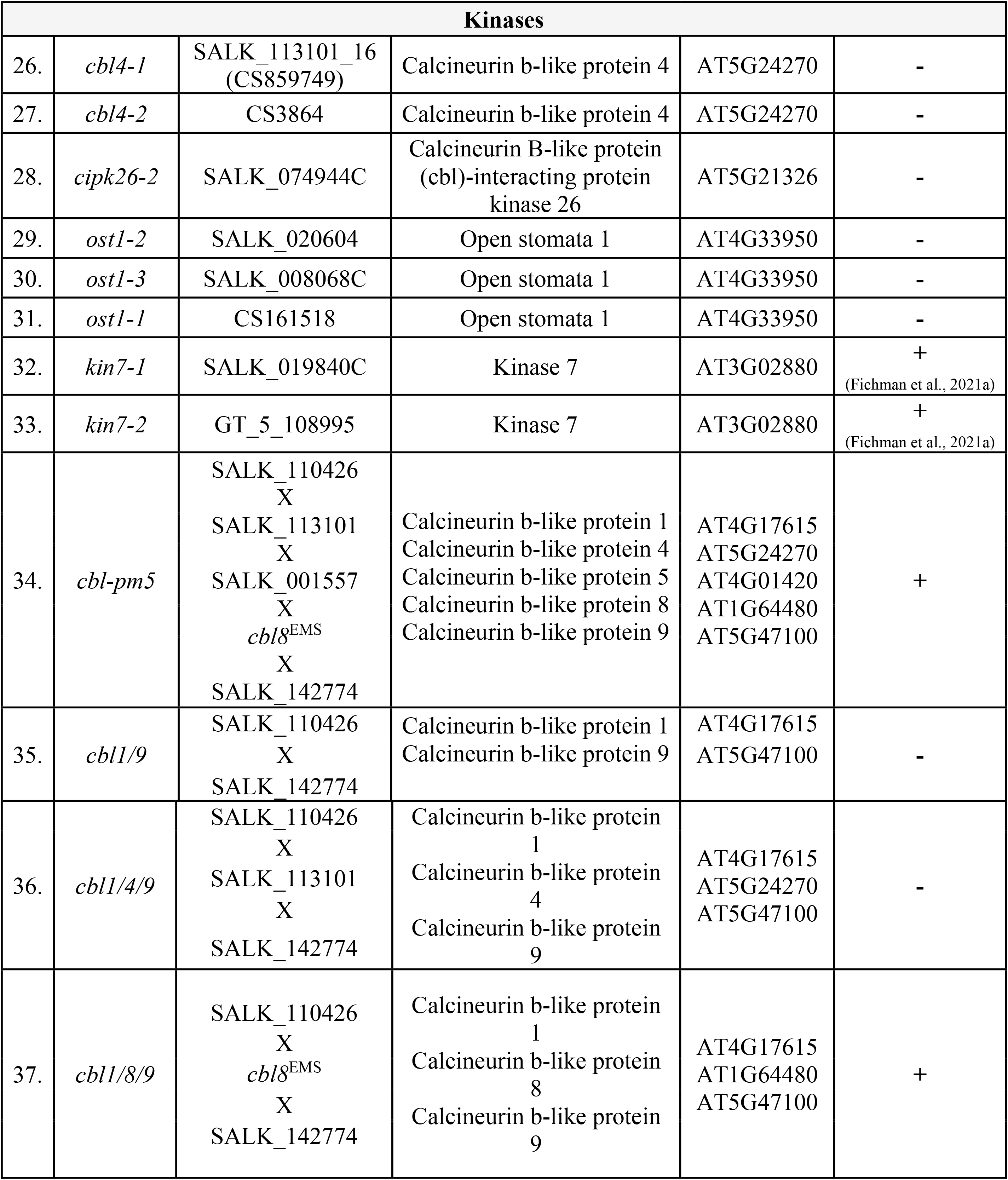
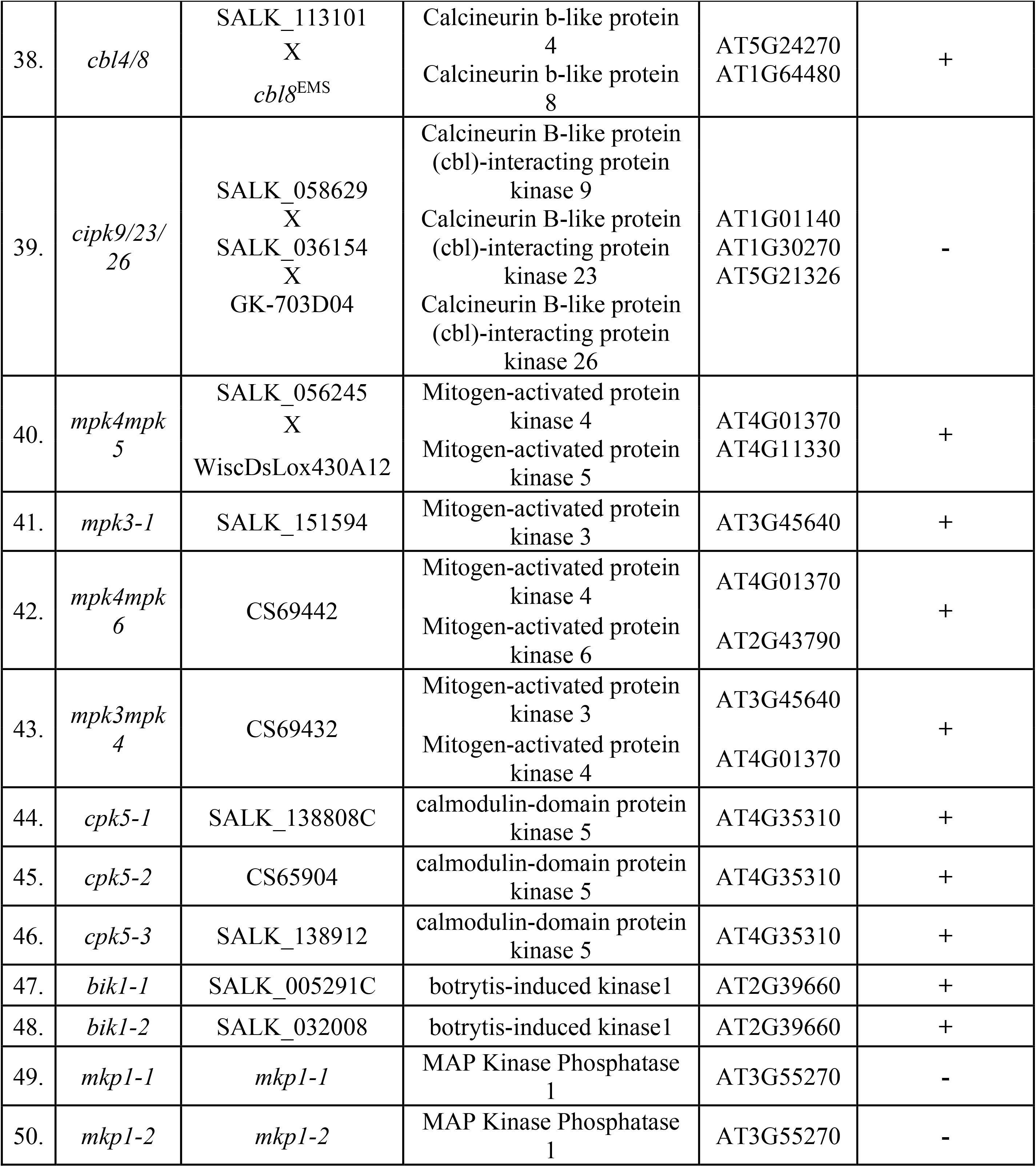
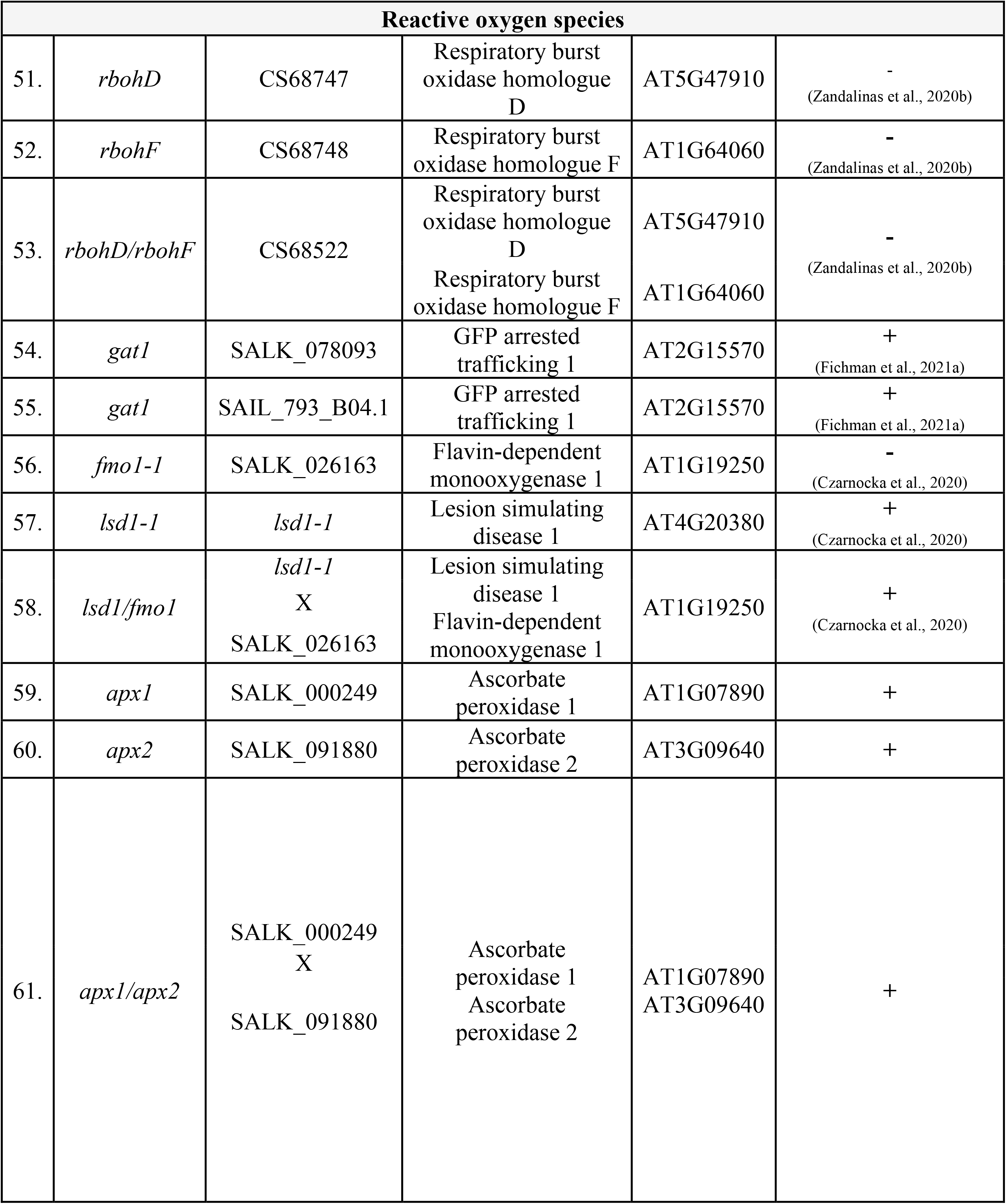
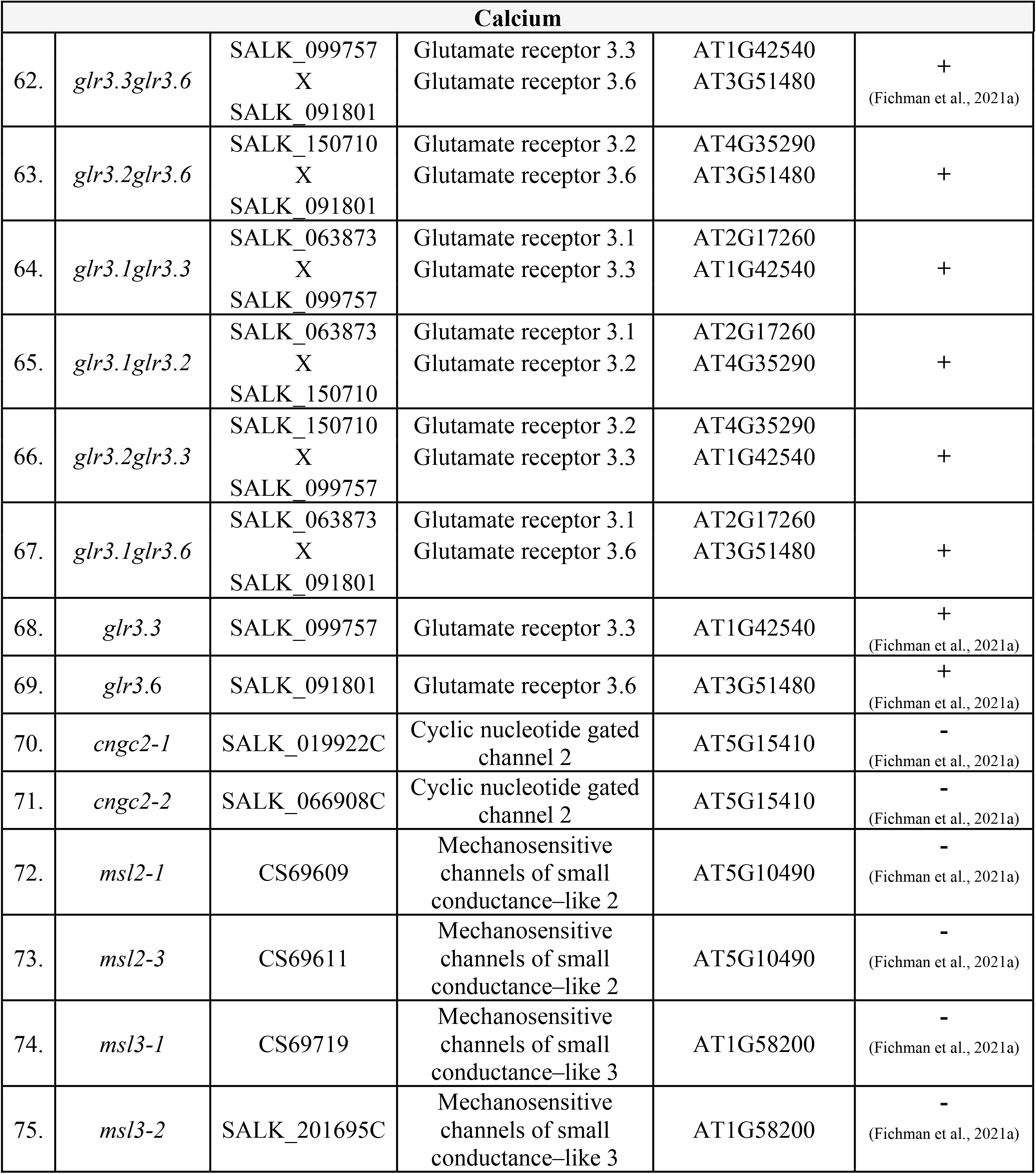
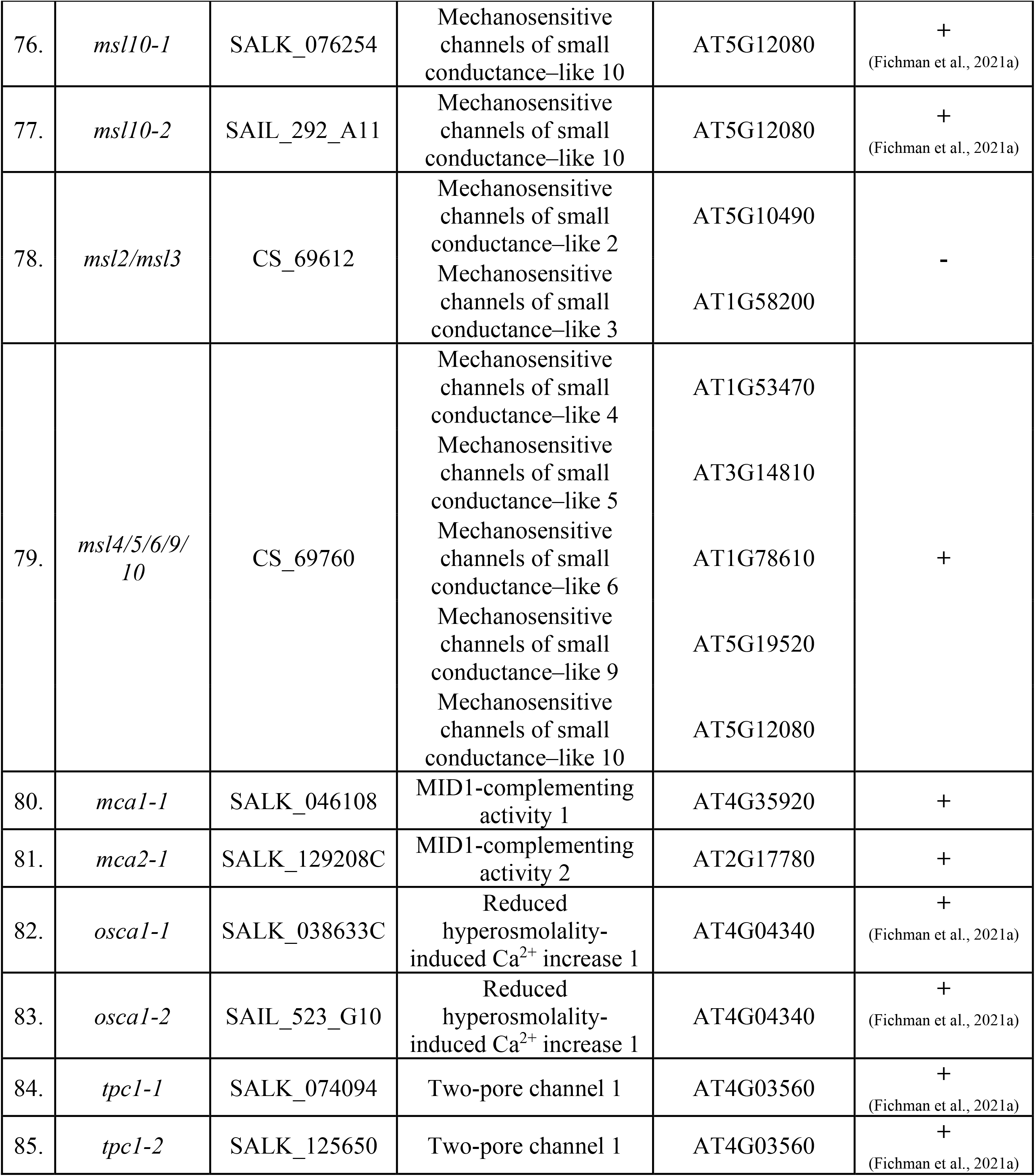
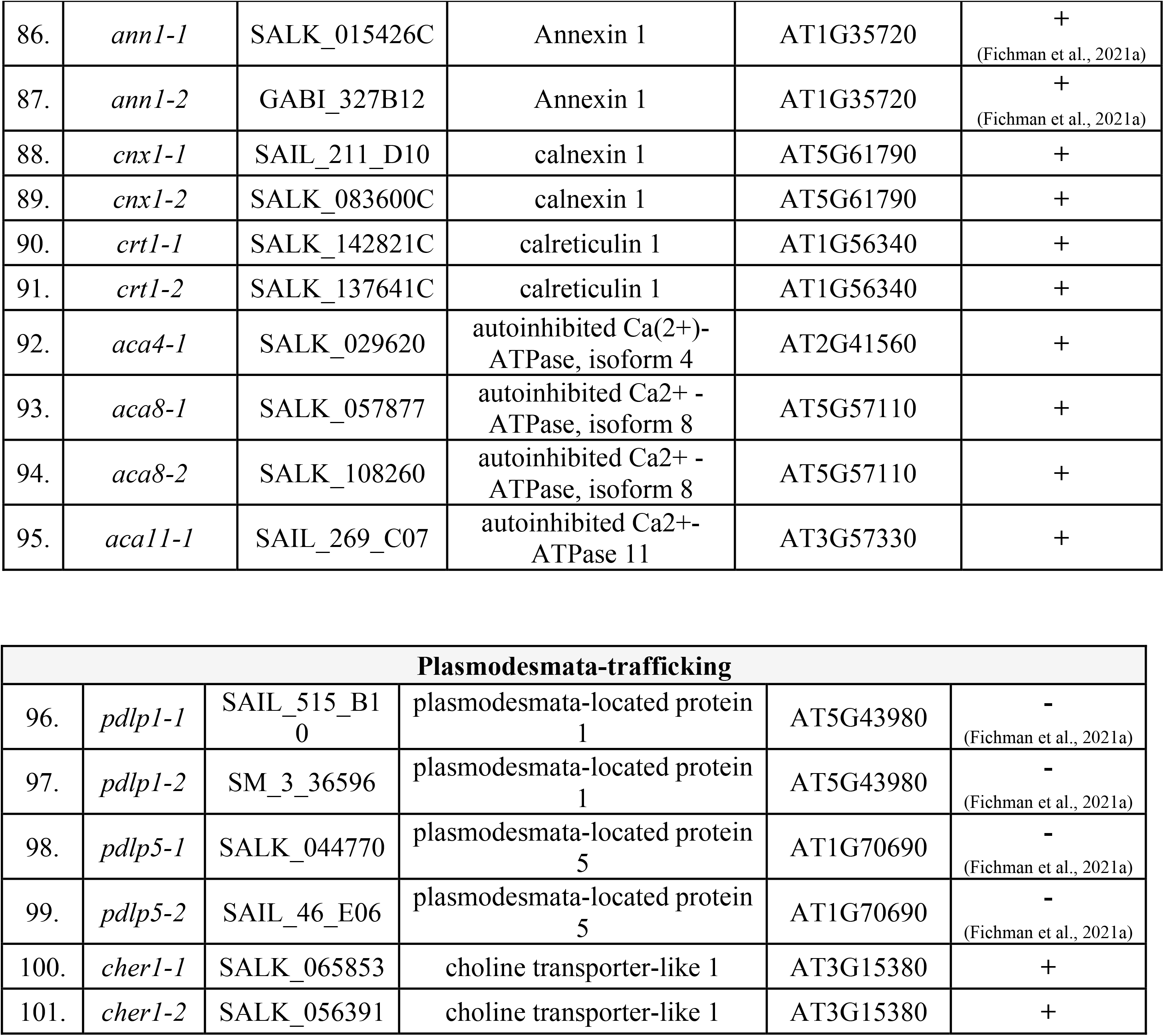
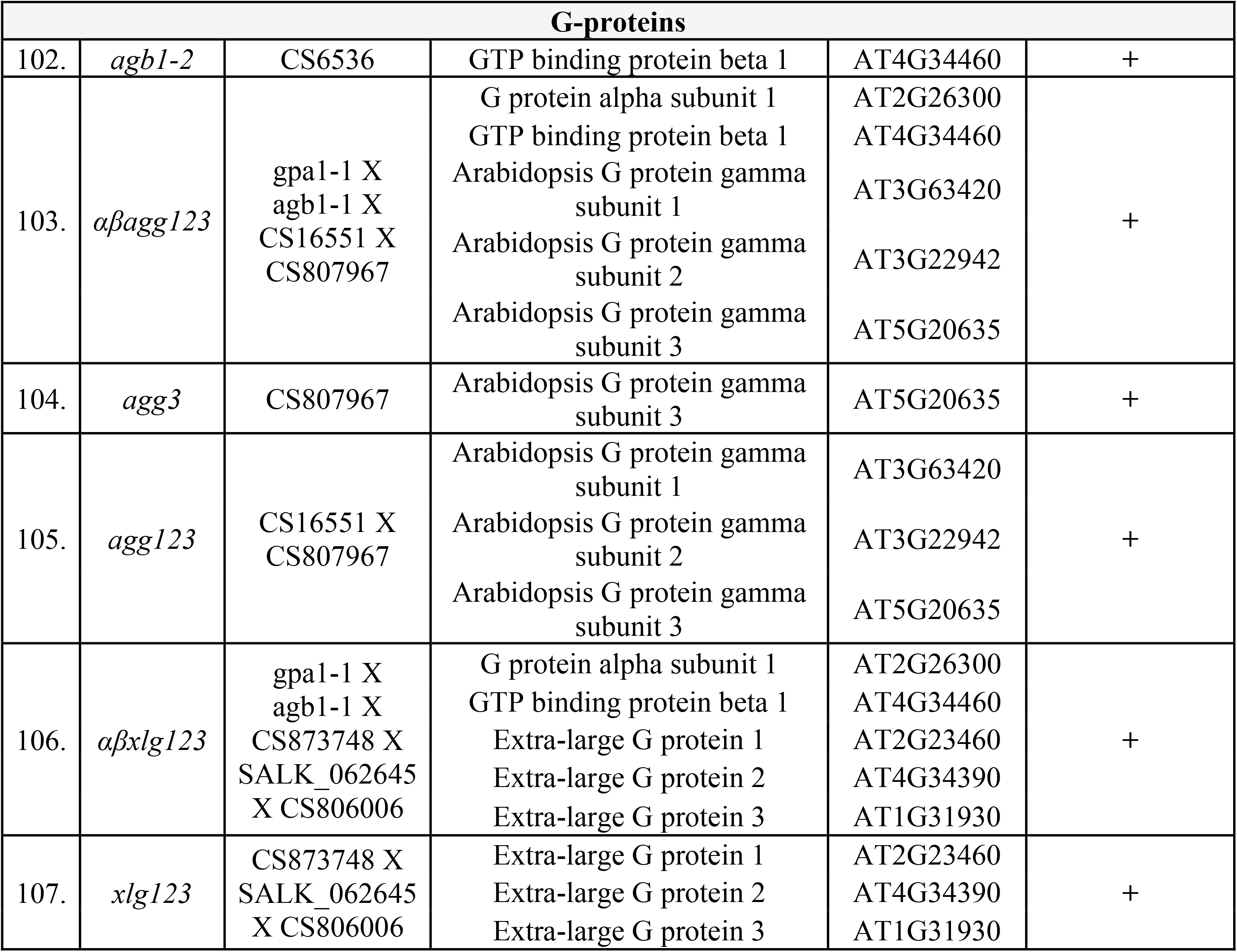
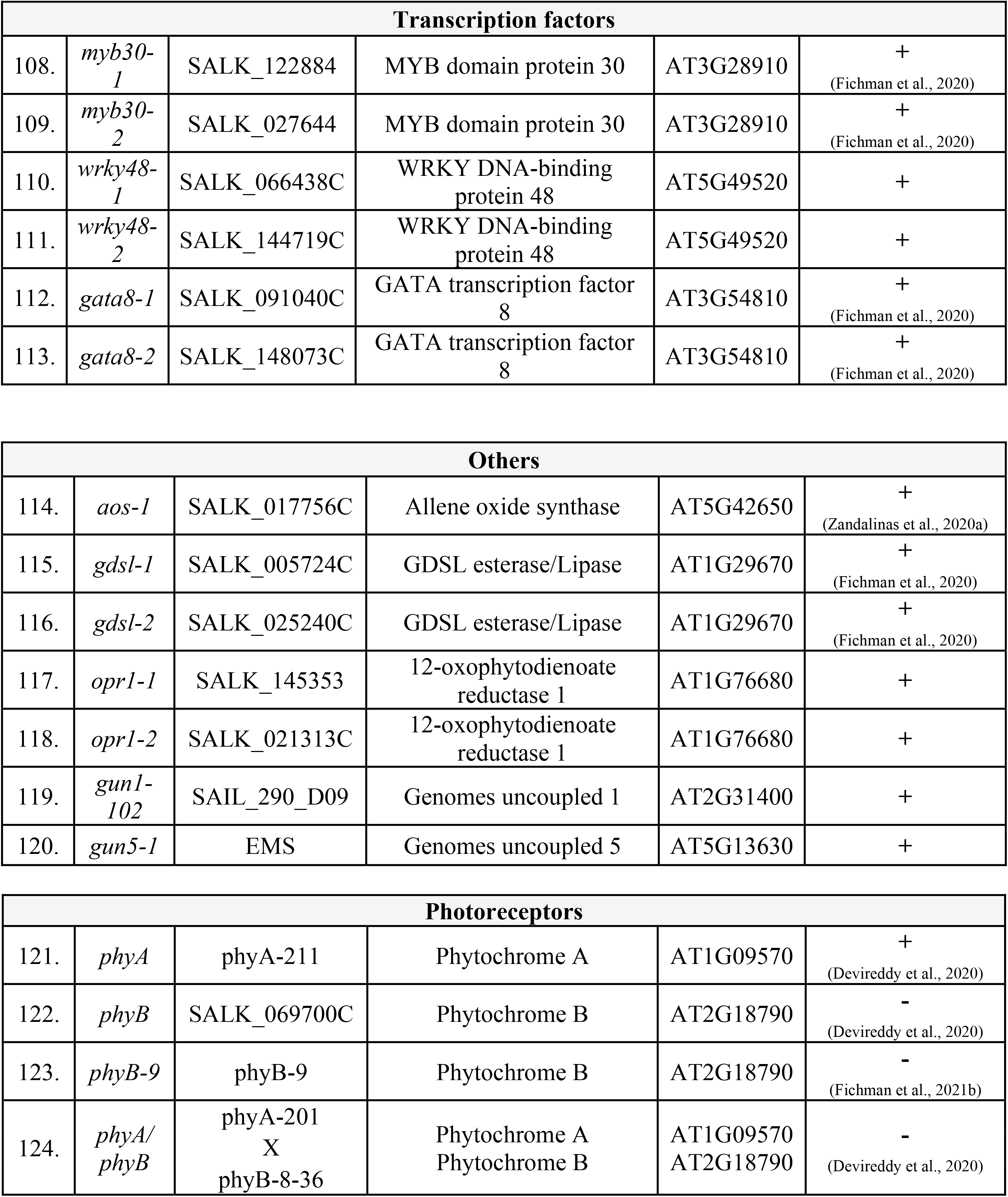
List of mutants that were screened for the presence or absence of the systemic ROS wave in response to a local highlight stress applied to a single leaf.

**Fig. S1.**
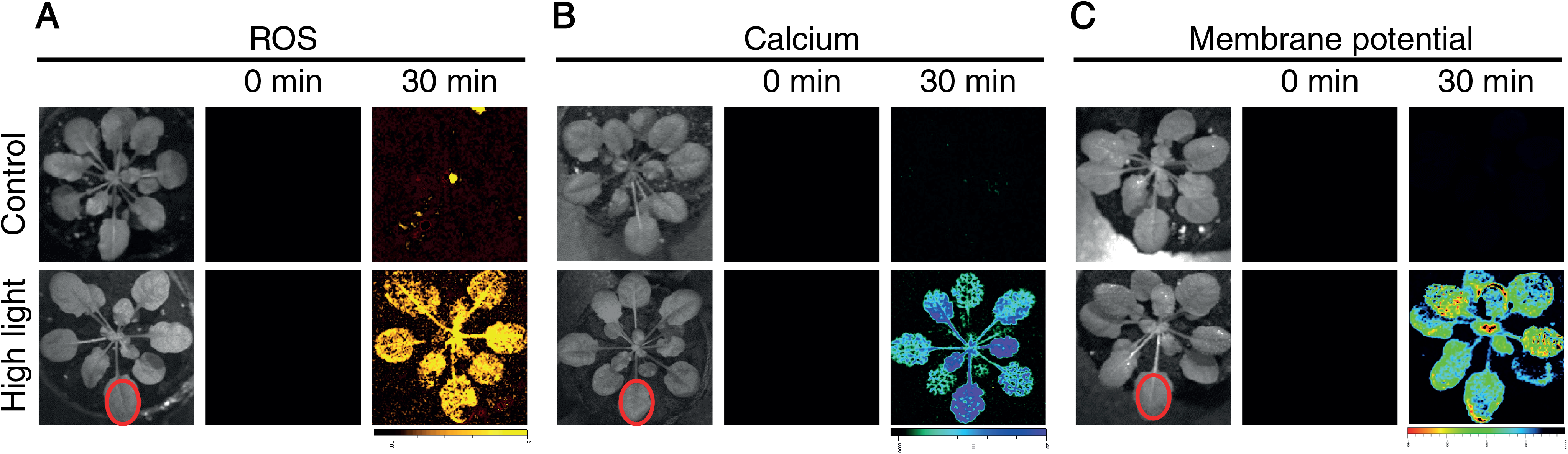
Imaging of ROS, calcium, and membrane potential in wild-type plants subjected to a HL stress treatment applied to a single leaf. *Arabidopsis* plants were untreated or subjected to a high light (HL) stress treatment applied to a single leaf (Local; indicated with a red circle), and ROS **(A)**, calcium **(B)**, or membrane potential **(C)** were imaged, using H_2_DCFDA, Fluo-4-AM, or DiBAC_4_(3), respectively, in whole plants (local and systemic tissues) as described in Fichman and Mittler (2021a), and the Methods section.

**Movie S1.** Live whole plant imaging of changes in cell-to-cell reactive oxygen species, calcium, and membrane potential signals in response to the application of high light stress to a single leaf (indicated by a white circle) of wild type and two independent mutants of HPCA1 (*hpca1-1*, *hpca1-2*). Note that although the detection of changes in cytosolic calcium levels precedes that of reactive oxygen species, cell-to-cell changes in calcium levels are dependent on reactive oxygen species sensing by HPCA1. By contrast, cell-to-cell changes in membrane potential do not require HPCA1.

## References

Aguirre, J., and Lambeth, J. D. (2010). Nox enzymes from fungus to fly to fish and what they tell us about Nox function in mammals. Free Radic. Biol. Med. 49:1342–1353.

Alonso, J. M., Stepanova, A. N., Leisse, T. J., Kim, C. J., Chen, H., Shinn, P., Stevenson, D. K., Zimmerman, J., Barajas, P., Cheuk, R., et al. (2003). Genome-wide insertional mutagenesis of Arabidopsis thaliana. Science 301:653–657.

Clough, S. J., and Bent, A. F. (1998). Floral dip: a simplified method for Agrobacterium -mediated transformation of Arabidopsis thaliana. Plant J. 16:735–743.

Dickinson, M. S., Lu, J., Gupta, M., Marten, I., Hedrich, R., and Stroud, R. M. (2022). Molecular basis of multistep voltage activation in plant two-pore channel 1. Proc. Natl. Acad. Sci. U.S.A. 119:e2110936119.

Drerup, M. M., Schlücking, K., Hashimoto, K., Manishankar, P., Steinhorst, L., Kuchitsu, K., and Kudla, J. (2013). The calcineurin B-like calcium sensors CBL1 and CBL9 together with their interacting protein kinase CIPK26 regulate the Arabidopsis NADPH oxidase RBOHF. Mol. Plant 6:559–569.

Evans, M. J., Choi, W.-G., Gilroy, S., and Morris, R. J. (2016). A ROS-assisted calcium wave dependent on the AtRBOHD NADPH oxidase and TPC1 cation channel propagates the systemic response to salt stress. Plant Physiol. 171:1771–1784.

Farmer, E. E., Gao, Y.-Q., Lenzoni, G., Wolfender, J.-L., and Wu, Q. (2020). Wound- and mechanostimulated electrical signals control hormone responses. New Phytol. 227:1037–1050.

Fichman, Y., and Mittler, R. (2020a). Rapid systemic signaling during abiotic and biotic stresses: is the ROS wave master of all trades? Plant J. 102:887–896.

Fichman, Y., and Mittler, R. (2020b). Noninvasive live ROS imaging of whole plants grown in soil. Trends Plant Sci. 25:1052–1053.

Fichman, Y., and Mittler, R. (2021a). Integration of electric, calcium, reactive oxygen species and hydraulic signals during rapid systemic signaling in plants. Plant J. 107:7–20.

Fichman, Y., and Mittler, R. (2021b). A systemic whole-plant change in redox levels accompanies the rapid systemic response to wounding. Plant Physiol. 186:4–8.

Fichman, Y., Miller, G., and Mittler, R. (2019). Whole-plant live imaging of reactive oxygen species. Mol. Plant 12:1203–1210.

Fichman, Y., Myers, R. J., Grant, D. G., and Mittler, R. (2021). Plasmodesmata-localized proteins and ROS orchestrate light-induced rapid systemic signaling in Arabidopsis. Sci. Signal. 14:eabf0322.

Gutteridge, J. M. C., and Halliwell, B. (2018). Mini-Review: Oxidative stress, redox stress or redox success? Biochem. Biophys. Res. Commun. 502:183–186.

Hörandl, E., and Speijer, D. (2018). How oxygen gave rise to eukaryotic sex. Proc. Royal Soc. B 285:20172706.

Iwashita, H., Castillo, E., Messina, M. S., Swanson, R. A., and Chang, C. J. (2021). A tandem activity-based sensing and labeling strategy enables imaging of transcellular hydrogen peroxide signaling. Proc. Natl. Acad. Sci. U.S.A. 118:e2018513118.

Jabłońska, J., and Tawfik, D. S. (2021). The evolution of oxygen-utilizing enzymes suggests early biosphere oxygenation. Nat Ecol Evol 5:442–448.

Johns, S., Hagihara, T., Toyota, M., and Gilroy, S. (2021). The fast and the furious: rapid long-range signaling in plants. Plant Physiol. 185:694–706.

Koncz, C., and Schell, J. (1986). The promoter of TL-DNA gene 5 controls the tissue-specific expression of chimaeric genes carried by a novel type of Agrobacterium binary vector. Molec Gen Genet 204:383–396.

Laohavisit, A., Wakatake, T., Ishihama, N., Mulvey, H., Takizawa, K., Suzuki, T., and Shirasu, K. (2020). Quinone perception in plants via leucine-rich-repeat receptor-like kinases. Nature 587:92–97.

Luan, S., and Wang, C. (2021). Calcium Signaling Mechanisms Across Kingdoms. Annu. Rev. Cell Dev. Biol. 37:311–340.

Lyzenga, W. J., Liu, H., Schofield, A., Muise-Hennessey, A., and Stone, S. L. (2013). Arabidopsis CIPK26 interacts with KEG, components of the ABA signalling network and is degraded by the ubiquitin–proteasome system. J. Exp.l Bot. 64:2779–2791.

Maurel, C., Tournaire-Roux, C., Verdoucq, L., and Santoni, V. (2021). Hormonal and environmental signaling pathways target membrane water transport. Plant Physiol. 187:2056–2070.

Miller, G., Schlauch, K., Tam, R., Cortes, D., Torres, M. A., Shulaev, V., Dangl, J. L., and Mittler, R. (2009). The plant NADPH oxidase RBOHD mediates rapid systemic signaling in response to diverse stimuli. Sci. Signal. 2:ra45–ra45.

Mittler, R. (2017). ROS are good. Trends Plant Sci. 22:11–19.

Mittler, R., Vanderauwera, S., Suzuki, N., Miller, G., Tognetti, V. B., Vandepoele, K., Gollery, M., Shulaev, V., and Van Breusegem, F. (2011). ROS signaling: the new wave? Trends Plant Sci. 16:300–309.

Mittler, R., Zandalinas, S. I., Fichman, Y., and Van Breusegem, F. (2022). ROS signalling in plant stress responses. Nat. Rev. Mol. Cell Biol. In press.

Mousavi, S. A. R., Chauvin, A., Pascaud, F., Kellenberger, S., and Farmer, E. E. (2013). GLUTAMATE RECEPTOR-LIKE genes mediate leaf-to-leaf wound signalling. Nature 500:422–426.

Nguyen, C. T., Kurenda, A., Stolz, S., Chételat, A., and Farmer, E. E. (2018). Identification of cell populations necessary for leaf-to-leaf electrical signaling in a wounded plant. Proc. Natl. Acad. Sci. U.S.A. 115:10178–10183.

Nühse, T. S., Bottrill, A. R., Jones, A. M. E., and Peck, S. C. (2007). Quantitative phosphoproteomic analysis of plasma membrane proteins reveals regulatory mechanisms of plant innate immune responses. Plant J. 51:931–940.

Ogasawara, Y., Kaya, H., Hiraoka, G., Yumoto, F., Kimura, S., Kadota, Y., Hishinuma, H., Senzaki, E., Yamagoe, S., Nagata, K., et al. (2008). Synergistic activation of the Arabidopsis NADPH oxidase AtrbohD by Ca2+ and phosphorylation. J. Biol. Chem. 283:8885–8892.

Razzell, W., Evans, I. R., Martin, P., and Wood, W. (2013). Calcium flashes orchestrate the wound inflammatory response through DUOX activation and hydrogen peroxide release. Curr. Biol. 23:424–429.

Rodrigues, O., Reshetnyak, G., Grondin, A., Saijo, Y., Leonhardt, N., Maurel, C., and Verdoucq, L. (2017). Aquaporins facilitate hydrogen peroxide entry into guard cells to mediate ABA- and pathogen-triggered stomatal closure. Proc Natl Acad Sci USA 114:9200–9205.

Schieber, M., and Chandel, N. S. (2014). ROS Function in Redox Signaling and Oxidative Stress. Curr. Biol. 24:R453–R462.

Shao, Q., Gao, Q., Lhamo, D., Zhang, H., and Luan, S. (2020). Two glutamate- and pH-regulated Ca2+ channels are required for systemic wound signaling in Arabidopsis. Sci. Signal. 13:aba1453.

Sies, H., and Jones, D. P. (2020). Reactive oxygen species (ROS) as pleiotropic physiological signalling agents. Nat Rev Mol Cell Biol 21:363–383.

Sirichandra, C., Gu, D., Hu, H.-C., Davanture, M., Lee, S., Djaoui, M., Valot, B., Zivy, M., Leung, J., Merlot, S., et al. (2009). Phosphorylation of the Arabidopsis AtrbohF NADPH oxidase by OST1 protein kinase. FEBS Letters 583:2982–2986.

Smirnoff, N., and Arnaud, D. (2019). Hydrogen peroxide metabolism and functions in plants. New Phytol. 221:1197–1214.

Szechynska-Hebda, M., Lewandowska, M., Witoń, D., Fichman, Y., Mittler, R., and Karpinski, S. (2022). Aboveground plant-to-plant electrical signaling mediates network acquired acclimation. Plant Cell In press.

Taverne, Y. J., Merkus, D., Bogers, A. J., Halliwell, B., Duncker, D. J., and Lyons, T. W. (2018). Reactive oxygen species: Radical factors in the evolution of animal life. BioEssays 40:1700158.

Torres, M. A., Dangl, J. L., and Jones, J. D. G. (2002). Arabidopsis gp91phox homologues AtrbohD and AtrbohF are required for accumulation of reactive oxygen intermediates in the plant defense response. Proc. Natl. Acad. Sci. U.S.A. 99:517–522.

Toyota, M., Spencer, D., Sawai-Toyota, S., Jiaqi, W., Zhang, T., Koo, A. J., Howe, G. A., and Gilroy, S. (2018). Glutamate triggers long-distance, calcium-based plant defense signaling. Science 361:1112–1115.

Wang, P., Hsu, C.-C., Du, Y., Zhu, P., Zhao, C., Fu, X., Zhang, C., Paez, J. S., Macho, A. P., Tao, W. A., et al. (2020). Mapping proteome-wide targets of protein kinases in plant stress responses. Proc. Natl. Acad. Sci. U.S.A. 117:3270–3280.

Waszczak, C., Carmody, M., and Kangasjärvi, J. (2018). Reactive oxygen species in plant signaling. Annu Rev Plant Biol 69:209–236.

Wu, F., Chi, Y., Jiang, Z., Xu, Y., Xie, L., Huang, F., Wan, D., Ni, J., Yuan, F., Wu, X., et al. (2020). Hydrogen peroxide sensor HPCA1 is an LRR receptor kinase in Arabidopsis. Nature 578:577–581.

Yang, Y., Zhang, C., Tang, R.-J., Xu, H.-X., Lan, W.-Z., Zhao, F., and Luan, S. (2019). Calcineurin B-like proteins CBL4 and CBL10 mediate two independent salt tolerance pathways in Arabidopsis. Int. J. Mol. Sci. 20:2421.

Zandalinas, S. I., Fichman, Y., Devireddy, A. R., Sengupta, S., Azad, R. K., and Mittler, R. (2020a). Systemic signaling during abiotic stress combination in plants. Proc. Natl. Acad. Sci. U.S.A. 117:13810–13820.

Zandalinas, S. I., Fichman, Y., and Mittler, R. (2020b). Vascular bundles mediate systemic reactive oxygen signaling during light stress. Plant Cell 32:3425–3435.

Zheng, H., Kim, J., Liew, M., Yan, J. K., Herrera, O., Bok, J. W., Kelleher, N. L., Keller, N. P., and Wang, Y. (2015). Redox metabolites signal Polymicrobial biofilm development via the NapA oxidative stress cascade in Aspergillus. Curr. Biol. 25:29–37.

Zhu, J.-K. (2016). Abiotic stress signaling and responses in plants. Cell 167:313–324.

## References

Czarnocka, W., Fichman, Y., Bernacki, M., Różańska, E., Sańko-Sawczenko, I., Mittler, R., and Karpiński, S. (2020). FMO1 is involved in excess light stress-induced signal transduction and cell death signaling. Cells 9:2163.

Devireddy, A. R., Liscum, E., and Mittler, R. (2020). Phytochrome B is required for systemic stomatal responses and reactive oxygen species signaling during light stress. Plant Physiol. 184:1563–1572.

Fichman, Y., Zandalinas, S. I., Sengupta, S., Burks, D., Myers, R. J., Azad, R. K., and Mittler, R. (2020). MYB30 orchestrates systemic reactive oxygen signaling and plant acclimation. Plant Physiol 184:666–675.

Fichman, Y., Myers, R. J., Grant, D. G., and Mittler, R. (2021a). Plasmodesmata-localized proteins and ROS orchestrate light-induced rapid systemic signaling in Arabidopsis. Sci. Signal. 14:eabf0322.

Fichman, Y., Xiong, H., Sengupta, S., Azad, R. K., Hibberd, J. M., Liscum, E., and Mittler, R. (2021b). Phytochrome B regulates reactive oxygen signaling during abiotic and biotic stress in plants. BioRxiv doi:10.1101/2021.11.29.470478.

